# Cmai: Predicting Antigen-Antibody Interactions from Massive Sequencing Data

**DOI:** 10.1101/2024.06.27.601035

**Authors:** Bing Song, Kaiwen Wang, Saiyang Na, Jia Yao, Farjana J. Fattah, Mitchell S. von Itzstein, Donghan M. Yang, Jialiang Liu, Yaming Xue, Chaoying Liang, Yuzhi Guo, Indu Raman, Chengsong Zhu, Jonathan E. Dowell, Jade Homsi, Sawsan Rashdan, Shengjie Yang, Mary E. Gwin, David Hsiehchen, Yvonne Gloria-McCutchen, Prithvi Raj, Xiaochen Bai, Jun Wang, Jose Conejo-Garcia, Yang Xie, David E. Gerber, Junzhou Huang, Tao Wang

## Abstract

The interaction between antigens and antibodies (B cell receptors, BCRs) is the key step underlying the function of the humoral immune system in cancers and other biological contexts. The capability to profile the landscape of antigen-binding affinities of a vast number of BCRs will provide a powerful tool to reveal novel insights at unprecedented levels and will yield powerful tools for translational development. However, current experimental approaches for detecting antibody-antigen interactions are costly and time-consuming, and can only achieve low-to-mid throughput. On the other hand, bioinformatics tools in the field of antibody informatics mostly focus on optimization of antibodies given known binding antigens, which is a very different research question and of limited scope. In this work, we developed an innovative Artificial Intelligence tool, Cmai, to address the prediction of binding between antibodies and antigens that can be scaled to high-throughput sequencing data. We devised a biomarker metric based on the output from Cmai applied to high-throughput BCR sequencing data to predict the landscape of the antigen-binding affinities of the BCRs. We found that extracellular antigens on malignant tumor cells are inducing B cell infiltrations, and the infiltrating B cells have a greater tendency to co-localize with tumor cells expressing these antigens. We further found that the abundance of tumor antigen-targeting antibodies is predictive of immune-checkpoint inhibitor (ICI) treatment response. On the other hand, we found that, during immune-related adverse events (irAEs) caused by ICI treatment, humoral immunity is preferentially responsive to intracellular antigens from the organs affected by the irAEs. Overall, we developed a powerful tool for elucidating the antigen binding landscape of BCR repertoires, and it shall also propell antibody-centric translational applications for cancers and other diseases.

## INTRODUCTION

B-cell receptors (BCRs) recognize diverse epitopes on antigen proteins and control the activation and maturation of B cells^1–4^. B cells with mature BCRs differentiate into plasma cells that secrete antibodies, which are the secreted forms of BCRs and carry out a variety of functions, such as neutralization of invading pathogens^5,6^. In addition to their key roles in infectious diseases and autoimmune diseases, recent studies have discovered curious parts that tumor-infiltrating B lymphocytes play in all stages of cancers, potentially in a BCR/antibody-dependent manner^7–11^. Given these, the capability to profile the binding affinities of any BCRs/antibodies in a repertoire against any antigens of interest can yield powerful insights into the function of the humoral immunity system in cancers and other biomedical contexts, and can lead to useful diagnostic biomarkers.

Traditional experimental approaches to identify antigen-antibody interactions are time-consuming and costly. The recent invention of Libra-seq^12,13^ has addressed this issue to some extent, which provides pairing information for thousands of BCRs against up to 20 antigens in a single experiment. 10X Genomics has also enabled capturing of the information of BCR-antigen binding affinity, through their UMI feature-barcoding method (Beam-B). Asrat *et al* have developed a similar technology called TRAPnSeq^14^. Related to these, Mohan *et al* developed PhIP-seq to map linear epitopes recognized by antibodies in a high-throughput manner^15^.

However, these experimental technologies are still limited in that they can only analyze a small set of given antigens, and can only scale to thousands of BCRs at a time. Besides, the time-consuming and costly aspects of these experiments are still not addressed. To achieve a landscape view of the binding affinities of BCR repertoires, we have to resort to bioinformatics-based predictions.

However, the field so far has mostly focused on machine learning-assisted optimization of antibodies, given the binding antigens^16–22^. This is a related but totally different scientific problem from *de novo* detection of antibody-antigen interactions. The optimization of antibodies focuses on modification of an antibody that is already known to bind a specific location on a known antigen. On the other hand, high-throughput detection of antibody-antigen interactions involves the consideration of many possible antigens and many possible antibodies, which can bind anywhere on the surface of the antigens. This is arguably a much more difficult scientific problem than antibody optimization, which has received little attention from the field yet. The newly released AlphaFold3 model^23^ is not applicable in this case, either, as AlphaFold3 predicts the multimer structure of an antibody and an antigen, assuming that they bind. This is different from the prediction of binding or non-binding of antigens and antibodies. AlphaFold3 also sets strict limits on how many AlphaFold3 jobs can be run each day by outside users, which makes it infeasible for detection of antigen-antibody interactions from massive sequencing data.

To fill this void, we developed a deep contrastive learning model, Cmai (short for Contrastive Modeling for Antigen-antibody Interactions), to predict the binding between antigens and BCRs/antibodies from BCR repertoire sequencing data. We systematically validated Cmai on independent antigen-BCR pairing data. We then devised a biomarker metric based on Cmai’s output to indicate the binding activities of the BCRs in any repertoire against the antigen(s) of interest. We showed that our approach can elucidate the roles of B cells and BCRs in physiologies of cancers. We envision our work will transform antibody-centric translational applications in the long run.

## RESULTS

### Cmai: Deep Learning the Binding of Antigens and Antibodies

The core Cmai model has three modules. First, we designed a set of auto-encoders for embedding antibody (BCR) sequences numerically so that they are manageable for downstream machine learning steps. Second, we created an antigen embedding module, based on RoseTTAFold^24^ as the backbone, for embedding the structural information of antigens. And finally, we created a deep contrastive learning module for predicting the interactions between antigens and antibodies (**Fig. 1(a)** and **Fig. S1**).

**Fig. 1.**
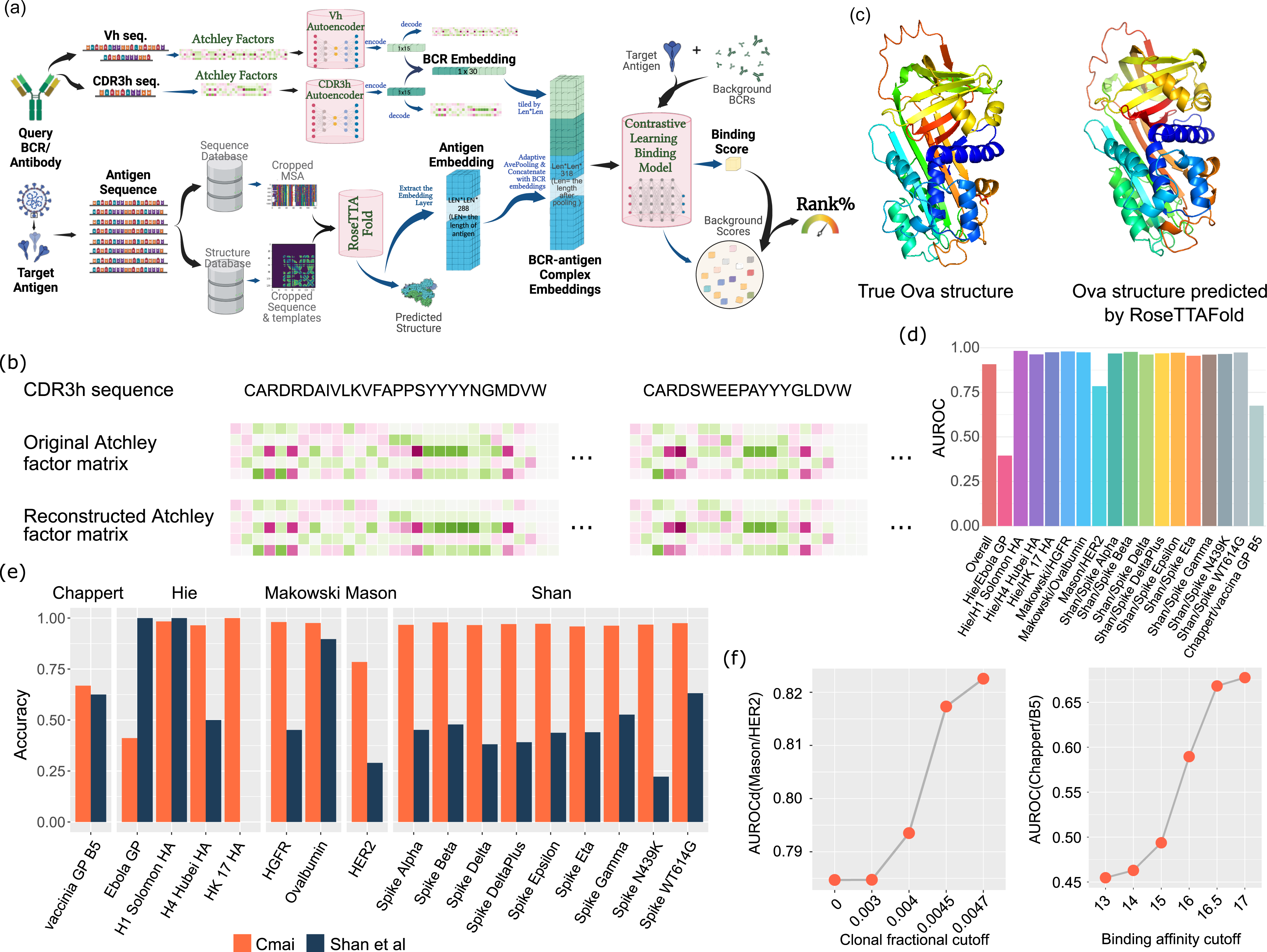
Cmai: deep learning the binding of antigens and antibodies. (a) Structure of the Cmai model. **(b)** The input and reconstructed Atchley factor matrices of two example BCR CDR3h sequences. (c) True OVA-alum structure and predicted OVA-alum structure. **(d)** AUROCs of Cmai on all antigens from all validation cohorts. (e) Accuracy of Cmai on antigens of all validation cohorts, where accuracy is defined as whether Cmai correctly predicted the true binding BCR to have lower rank% than the randomly sampled negative BCR, towards the antigen in the binding BCR-antigen pair. (f) The AUROCs on two validation cohorts increase as the stringency for defining true binding BCRs increases.

We collected 38,969 positive BCR-antigen pairing data from various sources and divided the pairing data into a training subset and an independent test subset (**File S1** and **Table S1**). The training dataset was further split into training and internal validation cohorts during the training Epochs. The training subset consists of 30003 positive binding pairs from 14 studies, while the test subset consists of 8,966 binding pairs from 5 studies. We have a total of 578 unique antigens in our collected pairing data (training+validation), which provides sufficient diversity of antigens for the training and validation of Cmai. The (2.5,97.5) % range of the antigen lengths is (99, 3,028), and the antigens are derived from various organisms (human, mouse, virus, bacteria, *etc*). The pairing data were derived from various biomedical contexts, such as cancers, infections, autoimmune diseases, *etc*. We ensured there is no overlap between the training and validation subsets. Then these data were used to train and validate Cmai. We employed a contrastive learning schema, where, for a positive binding antigen-BCR pair, we generated one negative BCR to pair with the same antigen through two distinct strategies: 1) “the binary negative BCR” for binary training phase: by mutating the positive BCR amino acid sequence randomly; 2) “the continuous negative BCR” for continuous training phase: by selecting a BCR with lower known binding affinity that also shares similar protein sequence. Details regarding the negative BCR generation are provided in the **method section**. We designed the loss function so that Cmai assigns a smaller loss for the positive binding pair, while assigns a larger loss for predicting the binding of the negative BCR towards the same antigen. Due to this design, at the prediction phase, we also employed a contrastive comparison schema for predicting the binding of a query antigen and a query BCR. In particular, we randomly sample 1 million BCR sequences to form a background distribution for the query BCR. The final binding prediction output is the rank percentile (rank%) of the predicted binding score of the query BCR-antigen pair among the scores output by the model for all the background BCR sequences against the same query antigen. A smaller rank% refers to a stronger predicted binding.

As one part of the input of Cmai, we developed a set of antibody embedding auto-encoders to numerically represent BCR/antibody sequences so that they can be understood by the machine learning model. In this part, we decided to focus on the heavy chains of the antibodies for the current Cmai model. This is due to concerns of the available pairing data’s sample sizes, where there is only the heavy chain information in many cases. Also, as has been shown before, the heavy chain accounts for the majority of the antigen binding specificity of the antibodies, compared with the light chains ^25,26^. We further divided the heavy chains into the Vh regions covering CDR1h and CDR2h, and the CDR3h regions, and designed two auto-encoders for each of Vh and CDR3h. Our rationale is that the Vh regions are longer but are known to be less important in determining antigen binding specificities of antibodies (sparser information), while the CDR3h regions are much shorter but are the key regions of the antibodies for determining binding specificities (denser information) ^25^. If we generate one single encoder for the whole heavy chain sequence, there is a risk that the encoder places more weight on the Vh regions simply due to the length of the Vh regions, while attending less to the CDR3h regions, which is suboptimal. Specifically, we created two variational auto-encoders (VAEs) for embedding the Vh and CDR3h regions, respectively. In essence, each amino acid of Vh and CDR3h is first embedded by Atchley factors^27^ as the input, and the output is a 15-neuron numerical vector for each of Vh and CD3h, taken from the bottleneck layers of the VAEs. We collected 7,743,963 BCR sequences (**File S1** and **Table S1**) to train these encoders. As we have shown in **Fig. 1(b)** for CDR3h and **Fig. S2** for Vh, the input Atchley factor matrices for CDR3h and Vh sequences are almost identical to the Atchley factor matrices reconstructed from the numerical embeddings of the auto-encoders (bottlenecks of the VAEs), which attests to the validity of our approach.

As the other part of the input, we also need to embed the antigen. We input the antigen sequences into RoseTTAFold and extracted the last layer in the RoseTTAFold model to form the antigen embeddings. The output embeddings for each antigen is in the format of a numeric matrix with the dimensions (LEN, LEN, 288), where LEN denotes the lengths of the antigen protein sequences and 288 refers to the number of feature channels. Moreover, the predicted protein structures are also saved as an intermediate model output (**Fig. 1(c)**), which facilitates diagnostics of the performances of the Cmai model in later stages. We note that many other protein structure prediction models such as AlphaFold^28^ and ESMFold^29^ may also be employed here for the purpose of the antigen embeddings. Each of these tools has its own advantages or disadvantages. We chose RoseTTAFold due to its embedding strategy that integrates various sources of information such as sequence, structure, and evolution, as well as a combination of other factors including computation speed, prediction accuracy, and ease of extraction of internal model layers.

At the final stage of the Cmai model, we concatenated the Vh and CDR3h encoders’ output into a 30-neuron numerical vector, as the overall BCRh embedding, and “inserted” the BCRh embedding at each position of the antigen embedding matrix to form a matrix of (LEN, LEN, 288+30) variables before the contrastive learning step. Our rationale is to mimic the natural process where the antibodies contact all parts of the antigen proteins and then determine the binding epitopes, *via* this *in silico* “insertion” operation.

### Cmai Achieves High Accuracy in Antigen-antibody Binding Prediction

We tested Cmai in our validation cohort, which consists of 8,966 positive binding pairs from 17 antigens from 5 cohorts (**Table S1**). For each positive binding pair, we created 10 negative pairs by randomly sampling 10 BCRs from our BCR pool and pairing these BCRs with the same antigen. The ultimate goal of Cmai is to identify antigen-BCR binding pairs from a pool of candidate antigens and a pool of candidate BCRs. Therefore, this method of forming the negative BCRs in the validation cohort is in the best alignment with this goal. However, one interesting observation we made is that the best training and validation outcomes result from negative BCR sets in our training data that were formed from BCRs similar to the positive binding BCRs in sequences (this is the case for both the binary and continuous negatives). One possible explanation is that this approach of forming similar positive-negative BCR sets feed BCRs of similar sequences to the Cmai model, so as to make it easier for the deep learning model to analyze and compare the two sequences and determine how the differences in sequences lead to differences in binding affinity. Otherwise, the model will be fed two strikingly different BCRs even during the warming up phase, which makes it difficult for Cmai to capture the key information that impacts binding.

We measured the performance of Cmai on these positive and negative pairs by the Area Under the Receiver Operating Characteristic (AUROC), as shown in **Fig. 1(d)**. For the majority of the antigens in the validation dataset, Cmai achieved AUROCs exceeding 0.95, with an average AUROC of 0.907. Cmai’s performance is the worst for the BCR-antigen pairing data for the Ebola Glycoprotein (GP) from Hie *et al*^21^. We conjecture that this might result from the poor protein structural prediction made by RoseTTAFold for this particular antigen. We compared the true Ebola GP structure from PDB (ID=6VKM) and predicted GP structure (**Fig. S3**). We indeed observed some discrepancies including the parts of the antigen that are close to the known antibody binding site, which might be the underlying reason for the lower AUROC.

We also tried to benchmark Cmai against competing software. However, we did not seem to find any other executable software tool that can predict the binding between antigens and antibodies in a high-throughput manner. Therefore, we executed the tool developed by Shan *et al*^19^, which is an antibody optimization tool by design. We created a benchmark dataset, where, for the same antibody-antigen positive binding pairs as used in **Fig. 1(d)**, we sampled only one negative BCR. We did this due to concern of computational speed as the Shan *et al* model requires multimer structure prediction for each pair, which is very slow. We deployed Cmai and Shan *et al* on this benchmark dataset, and measured the accuracy of prediction by whether the predicted binding is indeed stronger for the positive BCR than the negative BCR for all pairs. As is shown in **Fig. 1(e)**, Cmai out-performs the Shan *et al* model by a large margin (average accuracy across all cohorts is 0.91 for Cmai and 0.51 for Shan *et al*).

One caveat of our collected (or any) antibody-antigen pairing datasets is that there could be inaccuracies in the binding relationships, depending on the experimental technologies to derive the binding relationships. Fortunately, for some of these validation datasets, there are some recorded properties that are associated with the quality of the binding pairs. We took advantage of these metrics and defined more and more stringent subsets of antigen-BCR binding pairs for validation. We hypothesize that the prediction accuracy of Cmai should be higher in the higher quality subset. For example, in Mason *et al*^16^, we subset the HER2-binding BCRs by their observed clonal fractions, and we observed that Cmai’s accuracy is higher when measured with the more clonally expanded BCRs (**Fig. 1(f)**). The same is observed for Cmai’s prediction of the binding of BCRs towards vaccinia glycoprotein B5 (**Fig. 1(f)**), where the binding affinity is measured from the antigen-specific cell sorting by Chappert *et al*^16^. These results further validate the performance of Cmai.

### Cmai Recognizes Key Residues in Antigen-Antibody Binding Interfaces

To further examine whether Cmai is sensitive to small differences in the sequences of binding antigens and BCRs, we examined data from the Shan *et al* study^19^. In this study, we identified a total of 34 BCRs that are highly similar in their protein sequences and that are binding or not binding to several SARS-CoV-2 spike variants. We retained those BCRs that are sensitive to the different spike variants (**Fig. 2(a)**), meaning their coefficients of variation of log(IC50) for binding different antigens are smaller than −1. Then for each Spike variant, we examined the correlation between the predicted binding affinity (rank%, smaller means stronger binding) and log(IC50) (smaller means stronger binding) for the “sensitive” BCRs. As **Fig. 2(b)** shows, we indeed observed a positive correlation for each antigen, which suggests that Cmai can recognize small differences in antigens and BCRs that are likely in the key amino acid residues in the antigen-BCR binding interface.

**Fig. 2.**
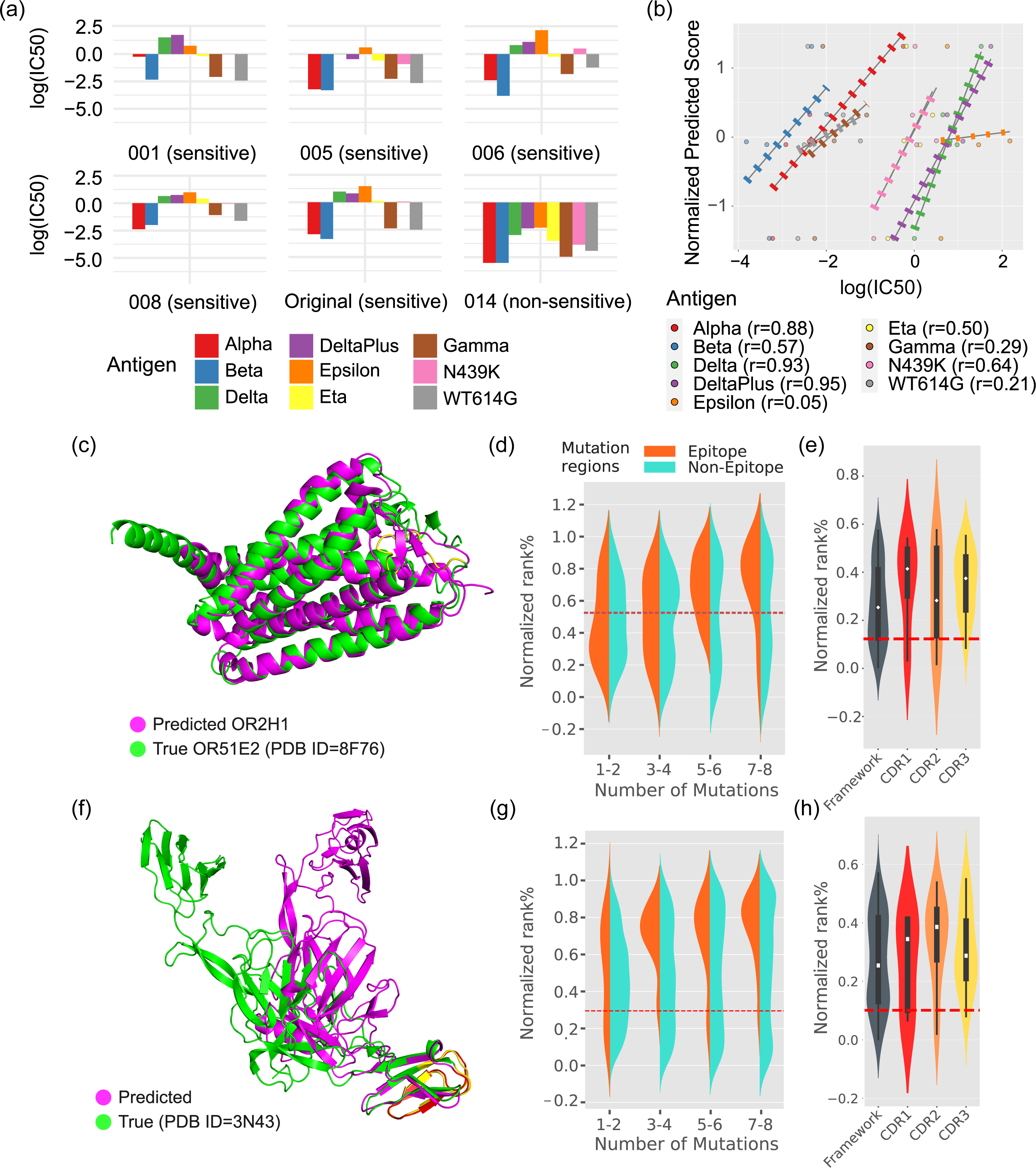
Cmai recognizes key residues in antigen-antibody binding interfaces. **(a)** In the Shan cohort, some BCRs uniformly bind all Spike protein variants (non-sensitive), while other BCRs only bind some of the variants (sensitive). Sensitive BCRs were defined as those with Coefficient of Variation of log(IC50) across all antigens smaller than −1. **(b)** The correlation between the predicted binding affinity (Y axis) and the true binding affinity (X axis) of the sensitive BCRs for each Spike variant. The binding affinity scores (rank%s) are normalized per Spike variant by mean-sd, for ease of visualization. (c) The true structure of human OR51E2 and the predicted structure of human OR2H1. The antibody binding epitope on OR2H1 is highlighted in yellow. (d) *In silico* alanine scanning results for OR2H1 by switching 1 to 8 contiguous amino acids to all alanines. The Y axis shows the predicted binding affinity by Cmai for all variant antigens. The rank%s are normalized among all variant antigens and shown on the Y axis for ease of visualization. On the X axis, the variant antigens are divided into “epitope” and “non-epitope” according to whether the center of the stretch of mutated amino acids falls within the binding epitope. (e) *In silico* alanine scanning results for all amino acids on the OR2H1-binding antibody heavy chain. One amino acid is switched to alanine at a time. **(f)** Predicted and true structures of the Chikungunya virus envelope protein. The antibody binding epitope on true and predicted antigens is highlighted in yellow and red, respectively. (g) *In silico* alanine scanning results for the Chikungunya virus envelope antigen protein. (h) *In silico* alanine scanning results for the Chikungunya virus envelope protein-binding antibody heavy chain.

Although explicit modeling of the interfaces or prediction of binding epitopes on the antigens is not the immediate goal of Cmai, we further confirmed the possibility that Cmai can recognize the key residues in the antigen-antibody binding interface in its internal calculation. To achieve this, we investigated a human OR2H1-BCR binding pair that we previously identified, for which we know the binding epitope: HQQIDDFLCEV ^30^. Cmai generated a prediction with a rank percentile of 3.6% for this pair, and the RoseTTAFold-predicted OR2H1 structure is highly similar to the true structure of OR51E2, the closest homolog with solved protein structure that we can find in the PDB database (**Fig. 2(c)**). We performed *in silico* alanine scanning in OR2H1 by switching contiguous segments (1-8 amino acids) of amino acids to alanines, which resulted in a total of 4,632 mutated OR2H1s. We predicted the binding of the BCR towards all these mutated antigens. As **Fig. 2(d)** shows, for antigens mutated in the binding epitope, there is a clear trend that predicted binding weakens as more and more switched alanines are introduced (T test P value=0.006 for rank% of 1-2 aa substitutions *vs.* 7-8 aa substitutions). But as a control, for the antigens that are not mutated in the epitope, the predicted binding has a much less obvious trend of weakening with increasing numbers of switched residues (T test P value=0.21 for rank% of 1-2 *vs.* 7-8 aa substitutions). On the other hand, we also performed alanine scanning for each amino acid in the BCR heavy chain. We observed that switching the amino acids to alanines will lead to weakened binding in all of the framework and CDR1-3 regions (Pval<1E-5 for all four regions), but the weakening of binding is stronger in the antigen-contacting CDR1-3 regions than in the framework regions (**Fig. 2(e))**, as expected.

We also investigated another known binding pair from SAbDab ^31^, which is a BCR bound to the glycoprotein of Chikungunya virus. As shown in **Fig. 2(f)**, RoseTTAFold successfully predicted the structure of this antigen protein. Note that the protein domain that contains the known binding epitope, K200G204G205S206E208K215V216I217N218N219, is connected to the rest of the antigen protein *via* flexible loops that can rotate freely in many angles. We performed alanine scanning up to 8 amino acids in the antigen, which resulted in 4,498 mutated antigens.

And we again observed that antigen variants mutated in the binding epitope have more weakened predicted binding with more alanines introduced (T test P value<1E-5 for rank% of 1-2 aa substitutions *vs.* 7-8 aa substitutions), while this trend is less clear for the variants that are not mutated in the binding epitope (T test P value=0.15 for rank% of 1-2 aa substitutions *vs.* 7-8 aa substitutions) (**Fig. 2(g)**). Similarly, we performed alanine scanning for each amino acid of the BCR heavy chain, and we again observed that alanine substitutions in the CDR regions lead to stronger weakened binding than substitutions in the framework regions (**Fig. 2(h)**).

### Targeting of Extracellular Tumor Antigens by Tumor Infiltrating B Cells

Next, we investigated whether Cmai could reveal any novel biological insights for the targeting of tumor antigens by tumor infiltrating B cells. We utilized data from the Cancer Genome Atlas (TCGA), which has collected over 20,000 tumor and normal samples from 33 cohorts, and performed RNA-sequencing and other types of genomics assays (**Fig. S4**). We extracted BCR sequences from the tumor RNA-sequencing data using mixcr^32^. A total of 6,002 samples with >20 unique BCR heavy chain sequences were retained. In parallel, we performed differential gene expression analyses between the normalized and log-transformed tumor and normal RNA-sequencing data and defined a set of 1,247 putative tumor antigens that fit the criteria of a minimum tumor expression cutoff and a minimum tumor/normal expression ratio cutoff in any cancer type, as laid out in our **method section**. We extracted the protein sequences for these putative tumor antigens from UniProt^33^. For each tumor sample, we predicted the binding of all the BCRs from that sample against the subset of the 1,247 tumor antigens that are expressed at a level that is higher than the minimum expression cutoff as mentioned above. Cmai was first used to predict the binding affinity between each patient sample’s BCRs against each putative tumor antigen. We then defined a biomarker metric, named “BCR binding score”, to measure whether the top expanded BCRs in each repertoire are enriched with BCRs that bind a certain tumor antigen of interest. The details of calculation are provided in the **method section**.

We investigated whether the expression levels of the tumor antigens are associated with the level of antibody responses. We first focused on the LUAD (Lung Adenocarcinoma) cohort. We calculated the expression of the putative tumor antigens found in LUAD with the expression averaged across all LUAD patients. We also calculated the average of the tumor antigen biomarker binding scores, across all LUAD patients for each tumor antigen. As **Fig. 3(a)** shows, there is indeed an overall positive correlation between the average tumor antigen expression and average tumor antigen binding strength by the LUAD patient BCRs. We then scaled this analysis to all cancer types (**Fig. 3(b)** and **Fig. S5(a)**), and confirmed that tumor antigens that are more highly expressed have induced overall stronger responses from the tumor infiltrating B cells, especially in the cancer cohorts with sufficient numbers of patients (**Fig. 3(b)**, N>100). In the chronic tumorigenesis processes, extracellular tumor antigens would have more exposure to the humoral immunity system, than intracellular antigens. We therefore hypothesize that this association will be stronger for extracellular antigens than for intracellular ones. We calculated the correlation between tumor antigen expression levels and tumor antigen binding scores, across extracellular and intracellular antigens. The definition of extracellular and intracellular antigens is provided in **Table S2**. As **Fig. 3(b)** and **Fig. S5(b)** show, the correlations are mostly positive for extracellular antigens as we suspected, while this is not observed for the intracellular antigens.

**Fig. 3.**
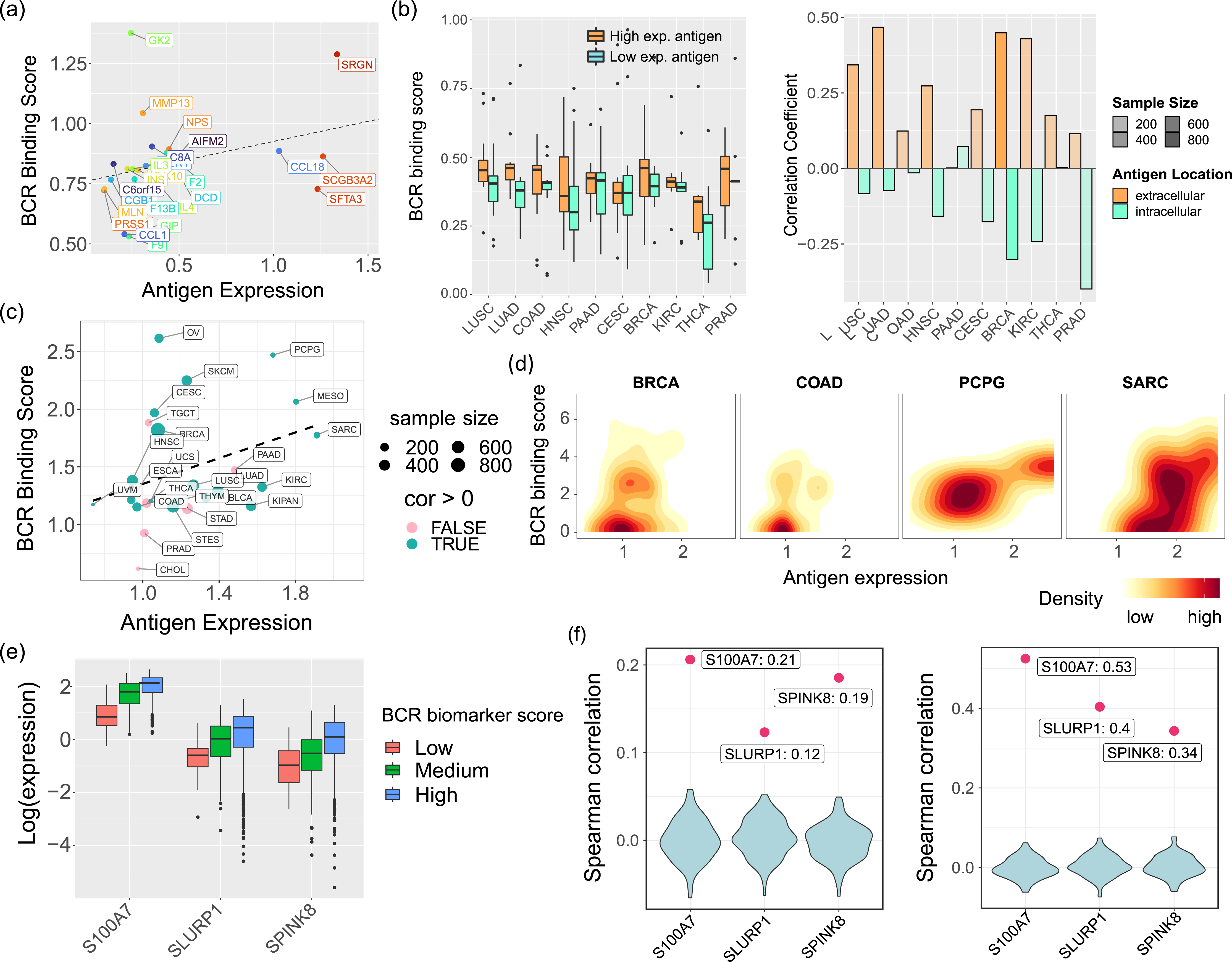
Cmai detects targeting of extracellular tumor antigens by tumor infiltrating B cells. **(a)** Scatterplot showing the average expression of tumor antigens with top expression (normalized and log-transformed expression>0.1) in the TCGA LUAD cohort (X axis) and the average BCR binding scores for these antigens across all LUAD patients. (b) Left: The average BCR binding scores are higher for tumor antigens that are more highly expressed, in each TCGA cohort. Right: The correlation between BCR binding scores and tumor antigen expression is higher for extracellular antigens than for intracellular antigens. In **(b)**, data are only shown for TCGA cohorts with >100 patients. (c) The average expression of *SRGN* and the average BCR binding score for *SRGN* across patients in each TCGA cohort. The pink and green color show whether the correlation between *SRGN* expression levels and *SRGN* binding scores is positive or negative across patients in each TCGA cohort. (d) Density plots showcasing the correlation between *SRGN* expression levels and SRGN binding scores in the patients from the TCGA BRCA, COAD, PCPG, and SARC cohorts. (e) Tumor sequencing spots in the c2 region with higher BCR binding scores for each of the three putative tumor antigens possess higher RNA expression levels of the tumor antigens. (f) Correlations between tumor antigen expression and BCR binding scores for the three putative tumor antigens and in both regions. The violin plots show the background distributions of the correlations that were generated by permuting the top binding BCRs for each tumor antigens.

In **Fig. 3(c)** (LUAD cohort), *SRGN* is the most highly expressed among the putative LUAD tumor antigens, which also possesses a top BCR binding score. SRGN, which encodes a proteoglycan involved in hematopoietic cell granule processes, has been linked to the regulation of granule-mediated apoptosis^34^. We next focused on *SRGN*, and showed its average gene expression levels and also the average BCR binding scores across all TCGA cohorts (**Fig. 3(c)**). Again, we observed that higher tumor *SRGN* gene expression is associated with stronger predicted binding by BCRs of the tumor infiltrating B cells. We looked within each cohort and examined the expression of *SRGN* and the SRGN binding score across all patients in each cohort (**Fig. 3(d)**). As **Fig. 3(CD)** show, the correlation coefficients are positive for most TCGA cohorts and the possibility of observing positive correlations is higher for cohorts with higher average expression of *SRGN*.

### Cmai Reveals BCR-dependent Co-localization of B cells and Tumor cells

Given that B cells infiltrate tumors dependent on BCR recognition of expressed tumor antigens, we hypothesize that tumor infiltrating B cells would tend to co-localize with the tumor cells that more highly express the antigens recognized by the B cells in the tumor microenvironment. Fortunately, Engblom *et al*^35^ recently incorporated the capability to sequence BCRs into Spatially Resolved Transcriptomics (SRT), which provided the data to address this question. Engblom *et al*^35^ applied their technology on a set of human breast cancer tissue samples in their work. We examined this dataset and focused on two samples with the best data quality (c2 and e2 regions, **Fig. S6(a)**).

We intersected the putative extracellular tumor antigens we defined for breast cancer above and the genes that are more highly expressed in the tumor cell regions over the stroma regions. This analysis yielded *S100A7* (expression pattern shown in **Fig. S6(b)**), *SLURP1*, and *SPINK8*, which we focused on in subsequent studies. We predicted, using Cmai, the binding between the protein sequences of these three putative antigens and all the BCRs found in these two regions. We calculated the BCR binding scores, as defined above, using the BCRs found in each sequencing spot and the corresponding Cmai predictions for each antigen.

We correlated the BCR binding scores with the expression of tumor antigens. We observed that the tumor antigen expression is higher in the tumor regions with higher antigen-specific BCR binding scores (**Fig. 3(e)**), while this is not observed when we divide the tumor regions by BCR clonal expansion sizes (**Fig. S6(c)**). As a background control, we created permutations by randomly permuting the order of BCRs in the calculation of BCR binding scores to form a background distribution, and the background BCR binding scores were correlated with expressions of tumor antigens. As is shown in **Fig. 3(f)**, the resulting correlations from the permutations are centered around 0, which suggests that the positive correlations we observed for the three putative tumor antigens are not a result of artifacts. Overall, our results reaffirm that B cells that target tumor antigens preferentially enter the tumor microenvironment and co-localize with tumor cells with high expression of the targeted antigens.

### The Cmai BCR Binding Scores Carry Prognostic and Predictive Information

As we have shown that B cells actively infiltrate and spatially distribute in tumors in a BCR dependent manner, we further evaluated whether the infiltration of B cells, measured by the BCR binding scores, carries any functional information. We first combined all TCGA cohorts and then averaged the BCR biomarker scores for the extracellular tumor antigens of each TCGA cancer type to serve as a measurement of the overall level of BCR-dependent tumor-specific B cell activities in each tumor sample. Then we performed Kaplan-Meier analyses by correlating this aggregate tumor antigen BCR binding score with the overall survival of the patients. In this initial trial when we consider all cancer types together, we did not observe a difference between the BCR score-high and BCR score-low patients (**Fig. 4(a)**). However, we performed this analysis again, but in each TCGA cancer type separately. Interestingly, there are only three cohorts for which the Kaplan-Meier analyses revealed a significant prognostic power for the BCR binding score, KIRC (Kidney renal clear cell carcinoma) (**Fig. 4(b)**), MESO (Mesothelioma) (**Fig. 4(c)**), and PAAD (Pancreatic adenocarcinoma) (**Fig. 4(d)**). And in all three cancer types, higher BCR binding scores were found to be predictive of poor overall survival for the patients. Then we combined these three cohorts and performed a multivariate correction for the confounding factors of race, gender and age, *via* the CoxPH analyses (**Fig. 4(e)**. The association between higher BCR binding score and poor prognosis is still statistically significant, after this adjustment. We also performed this analysis with the binding scores calculated for the extracellular and intracellular antigens separately and made similar observations (**Fig. S7**). Prior literature reported both tumor promoting and inhibiting roles of B cells^36,37^ in various contexts, which might explain the lack of a clear association between the BCR binding scores and prognosis across all TCGA cohorts. However, our analyses suggest that, for some particular cancer types like KIRG, MESO and PAAD, the BCR binding scores are indicative of poor prognosis and tumor infiltrating B cells likely have played dominant tumor promoting roles.

**Fig. 4.**
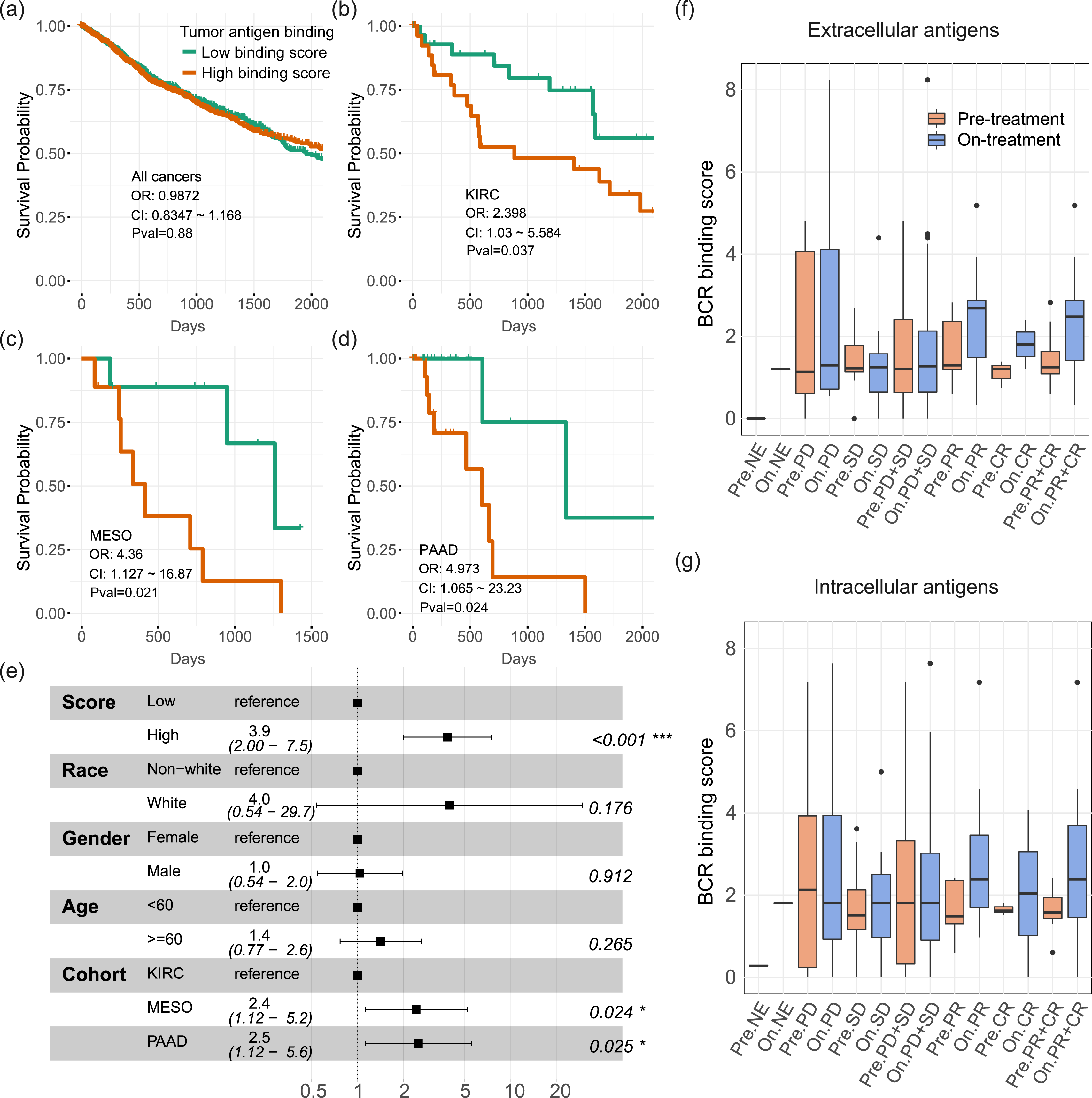
The Cmai BCR binding scores carry prognostic and predictive information. **(A-D)** The prognostic values of the Cmai binding scores. Patients were dichotomized based on median Cmai BCR binding scores in each cohort. (a) All TCGA patients; **(b)** TCGA KIRC patients; **(c)** TCGA MESO patients; and (d) TCGA PAAD patients. (e) Forest plot showing the results of the multivariate correction for the prognostic effects of the Cmai BCR binding scores, in the KIRC, MESO and PAAD patients combined. **(F,G)** The BCR binding scores in anti-PD1-treated patients, in each response category and before/after treatment. BCR binding scores are calculated for (f) extracellular and (g) intracellular tumor antigens, respectively.

We also evaluated whether the BCR binding scores are associated with treatment responsiveness to anti-PD1 treatment. Although *PD1* has been mostly studied for their effect on T cells in the context of immunotherapies, B cells also express *PD1*^38,39^ and whether B cells will be affected by PD1 blockade is not clear yet. We investigated an anti-PD1 treatment cohort from Riaz *et al*^40^, with a total of 68 melanoma patients on nivolumab treatment. 109 pre- and on-treatment samples were collected and RNA sequencing was performed. We extracted the BCR sequences from the RNA sequencing data using mixcr. And we examined the binding of the BCR repertoire from each patient towards the melanoma tumor antigens that we defined above and that were expressed higher than the minimum expression cutoff in each patient, as defined above. We calculated the tumor antigen biomarker scores for these antigens. We first focused on the extracellular antigens (**Fig. 4(f)**). In the pre-treatment samples, we did not observe any difference in the tumor antigen binding scores in the responders (complete response, CR; partial response, PR) *vs.* non-responders (stable disease, SD; progressive disease, PD). But strikingly, in the on-treatment samples, the tumor antigen binding scores increased significantly in the responders compared with the pre-treatment samples, while the scores stayed stable in the nonresponders.

For the intracellular antigens (**Fig. 4(g)**), we also observed the same trend, although with less extent of statistical significance (T test Pval for extracellular antigens: 0.000865; Pval for intracellular antigens: 0.00946). These results suggest that B cells are also effectively mobilized during anti-PD1 blockade. Further research is warranted by these results to investigate if B cells could actively affect responses to anti-PD1 and other immune checkpoint inhibitors, or if their activation is just a byproduct of immunotherapy treatment.

### Cmai Interprets Autoantibody Dynamics during Toxicity Response to Checkpoint Blockade

Having established the performance of Cmai, we next demonstrated whether Cmai could reveal novel biological insights from the landscape of antigen-antibody interactions in a repertoire. We first studied our cohort of immune checkpoint inhibitor (ICI)-treated patients collected from UT Southwestern Medical Center. For these patients, we have captured comprehensive clinical records including immune-related adverse events (irAEs), autoimmune toxicities that occur during ICI treatment. irAEs convey substantial morbidity, incur considerable costs, and in some cases may preclude further use of ICI drugs. We have a total of 113 patients treated by ICIs (anti-PD1/-PDL1/-CTLA4, mono- or dual-therapy), from which we collected blood draws for a total of 256 samples. Patient demographic and clinical characteristics are summarized in **Table S3**. The blood collections happened at baseline, times of irAE occurrences and several pre-schedule time points, although not all patients completed blood collections at all time points due to logistics reasons. We have also performed bulk RNA-sequencing and auto-antibody profiling (see **method section**) from these biospecimens. We extracted BCR sequences from the bulk RNA-sequencing data using mixcr^32^.

We first examined the auto-antibody data. There are 11 auto-antigens for which IgG data points were collected for the majority of all our samples (>190 samples). We took advantage of these data to first validate whether Cmai can correctly profile the reactivities of the patient BCR repertoire in the blood against auto-antigens. We predicted the binding between these BCRs and each auto-antigen, and calculated the BCR binding score as mentioned above. As **Fig. 5(a)** shows, there is a high degree of concordance between the predicted BCR binding scores and the measured autoantibody levels across patient samples for these 11 auto-antigens.

**Fig. 5.**
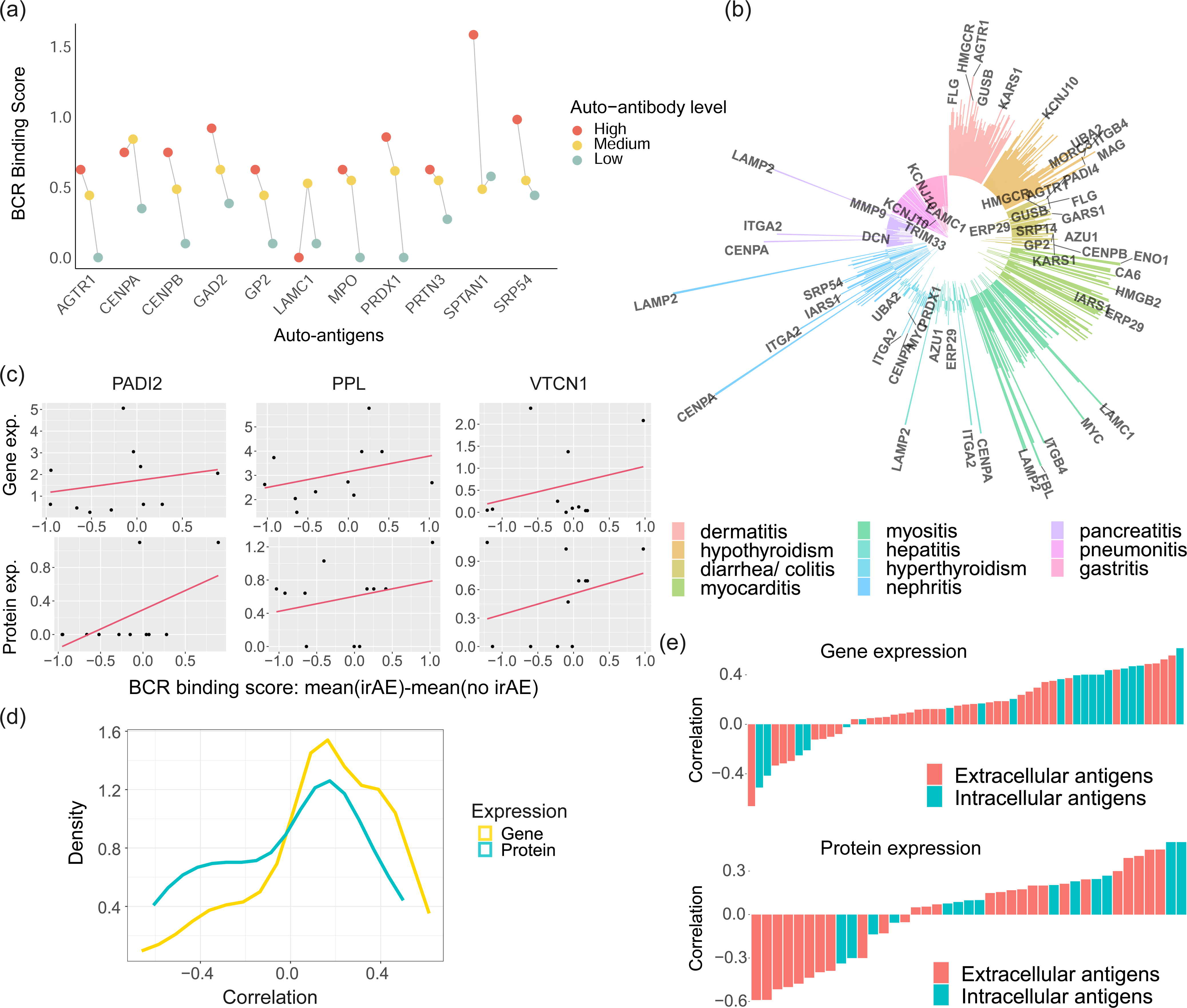
Cmai interprets autoantibody dynamics during toxicity response to checkpoint blockade. **(a)** The correlation between truly measured auto-antibody levels and the predicted auto-antibody levels by Cmai for the 11 auto-antigens for which we have sufficient auto-antibody data points. **(b)** Summary of the changes in the predicted auto-antibody levels in patient samples with irAEs over patient samples without irAEs, across all captured irAEs and all auto-antigens. Spike pointing outwards means the predicted auto-antibody level is higher in the irAE+ samples, while spikes pointing inwards mean the predicted auto-antibody level is lower in the irAE+ samples. (c) Scatterplots showing the correlation between the RNA and protein expression levels of three example auto-antigens in the organs affected by the irAEs *vs.* the elevation of the predicted auto-antibody levels when the irAEs affecting each organ happen. (d) Density plots showing the correlations between the BCR binding scores differences (irAE-no irAE) for each irAE and the gene/protein expressions of the organ affected by that irAE, for all auto-antigens. (e) Waterfall plots showing the distributions of the correlations, as calculated in panel (d), for all auto-antigens studied, based on either RNA expression or protein expression. Extracellular and intracellular antigens were separately color-coded.

We next correlated our predictions with occurrences of irAEs. For each irAE and each patient sample (blood collection time), we defined the patient sample as being associated with that irAE, if an irAE event has happened within (−30, 60) days of the corresponding blood collection time, and *vice versa*. We included a total of 63 commonly accepted auto-antigens (same as those on our auto-antibdoy testing panel, see **method section** and **Table S4**), as the prediction of binding can be performed as long as the antigen sequence can be identified. We calculated the same BCR binding score for each of these antigens across all patient samples. We observed that the predicted auto-antibody binding strengths to the auto-antigens mostly increased in the irAE-positive samples, compared with the irAE-negative samples, for dermatitis, diarrhea/colitis, myositis, myocarditis, and hypothyroidism (**Fig. 5(b)**). For pancreatitis, pneumonitis, gastritis, the autoantibody levels mostly decreased in the irAE-positive samples.

We hypothesize that one way, in which autoantibodies and autoantigens participate in the biological processes of irAEs, is when irAEs damage the healthy organs and release the auto-antigens into blood circulation. These auto-antigens will be recognized by the humoral immunity system and auto-antibodies will be generated against these antigens. If this is correct, the organ-specific expression of the auto-antigens would show an overall positive correlation with the extent of the elevations of the predicted auto-antibody binding scores, as the release of these auto-antigens stimulated the auto-antibody responses. To prove this, we first obtained the RNA gene expressions of these auto-antigens from the GTEx database^41^. We averaged the expression across all cell types that are found in each organ, to obtain the organ-specific gene expression for these auto-antigens (**Fig. S8**). Then we correlated the organ-specific auto-antigen expression with the extent of elevations of the auto-antibodies when the irAE affecting each organ happens, for each auto-antigen (**Fig. 5(c)**). As we expected, these correlation coefficients are indeed mostly positive (**Fig. 5(d)**, 76% of all antigens showed positive correlation, Pval of binomial test=2.7E-5). Furthermore, we hypothesize that this effect is likely to be more obvious when we focus on intracellular antigens rather than extracellular antigens. This is because the humoral immunity is likely to respond stronger to the transient release of the intracellular antigens upon irAEs, which are originally inaccessible to the humoral immunity system. As **Fig. 5(e)** shows, intracellular auto-antigens indeed possess correlation coefficients that are more positive than extracellular auto-antigens (median cor. of intracellular antigens=0.36, median cor. of extracellular antigens=0.12, KS test Pval=0.03).

We also examined protein expression data from Human Protein Atlas (HPA)^42^. We performed the same analyses (**Fig. 5(d)-(e)** and **Fig. S8**), and also made similar observations. Overall, Cmai confirmed our hypothesis that one of the major determinants of autoantibody levels during irAEs is the amount of released auto-antigens from the damaged organs that are recognized by the humoral immunity system.

## DISCUSSION

In this work, we developed an AI model to predict the binding between antibodies/BCRs and antigens in a high-throughput manner, and devised a biomarker approach based on this model to indicate the reactivity level of a BCR repertoire against any antigens of interest. This outcome is a result of our novel and delicate machine learning designs, but is also attributed to the timely progress in the field that have significantly enhanced the feasibility of our pursuit. First, works such as AlphaFold^23,28^, RosettaFold^24^, ESMFold^29^, *etc*, have enabled accurate prediction of protein tertiary structures based on amino acid sequences for many proteins, which can be leveraged to assist the prediction of antibody-antigen interaction. Second, the development of high-throughput sequencing-based technologies such as Libra-Seq, Beam-B and TRAPnSeq has propelled the accumulation of large amounts of antibody-antigen pairing data that would be necessary to train such AI models for the binding prediction tasks.

One important utility of our BCR-antigen binding prediction model is to employ it as a building block of BCR-based biomarkers against any antigens of interest. As long as the BCRs have been sequenced, this biomarker can be constructed for predicting the activity of the BCRs against any antigens without extra cost nor need to re-access the biospecimens again for experimental manipulations. Such biomarkers can be deployed to elucidate the relevance of the humoral immunity system in cancers and other human diseases, and can also be deployed for diagnosis and prognosis purposes. In the application of antibody optimization, experimental approaches are already very mature and AI-for-drug approaches for addressing this problem have a relatively smaller margin of advantage. In contrast, in the case of BCR-based biomarkers, which is enabled by Cmai, experimental approaches are much less feasible and less powerful. Cmai, which can identify antigen-antibody interactions from massive sequencing data, presents a significant advance addressing a need that has not been met yet.

One important design decision for Cmai is that we leveraged RoseTTAFold for modeling the structures of antigen proteins, but not for the antibodies. This is because these popular folding algorithms have been proved to be effective for predictions of many “general” proteins. These proteins commonly have many homologs, and these folding algorithms can more accurately predict their structures *via* evolutionary relationships and borrowing of information of solved structures through homologous relationships. But antibodies are a special class of proteins and there are not many solved structures yet, compared to what is necessary for training a very good antibody structure prediction model. Besides, the sequences of antibodies are flexible and not conserved, due to the abundance of somatic hypermutations that happen to the antibodies, especially in the most critical CDR regions. The CDR regions are also disordered loops without fixed structures. Therefore, we cannot deploy these popular folding algorithms to model antibodies. But rather, we developed our own BCR auto-encoders to numerically represent the protein sequences of antibodies for downstream machine learning purposes.

One feature that is likely important to consider in the future iterations of our work is the capability to identify the binding epitopes on the antigens. Our analyses suggest that the current Cmai model has implicitly considered the binding epitopes when predicting the binding between antigens and antibodies. With the accumulation of data generated from high-throughput technologies that can map the binding epitopes of the antigens bound by the antibodies, such as PhIP-Seq^15^, AI prediction of antibody-binding epitopes on the antigens will likely become possible in the future. Note this is different from and more difficult than prior works that just predict the possibility of epitopes on antigens without knowing the binding antibodies ^4,43,44^. When this advance is achieved, the biomarker metric that we have built on top of Cmai will readily become capable of indicating epitope-specific BCR reactivities, rather than overall antigen-specific BCR reactivities, offering a much higher level of resolution.

Another aspect of antigen-antibody binding to consider is that the isotype of antibodies could yield different biological functions. Biswas *et al*^45^ showed that all epithelial cancers express *PIGR* and internalize dimeric IgA, but not IgG, and generated a dimeric IgA that targets the intracellular antigen of *KRAS*^G12D^. This feature makes the profiling of antibody-antigen interactions more complicated than the prediction of binding between T cell receptors and antigens^46–51^, where there is no class switching involved. In our future work, we will incorporate the information of class switching, either as part of the Cmai model and the BCR binding metric, or as a confounding factor to adjust for in downstream analyses, as appropriate.

In balance, we have developed a powerful toolkit for profiling the landscape of antigen-binding affinities of massive BCR sequencing data, which will greatly facilitate both basic science research and personalized medicines where B cells, BCRs and antibodies are involved.

## MATERIALS AND METHODS

### Cmai model structure

The Vh and CDR3h sequences of BCR heavy chains are first embedded into Atchley factor matrices by replacing each amino acid with the 5 Atchley factors. Then variational auto-encoders were deployed for the Vh and CDR3h Atchley factor matrices, which consist of a hybrid architecture that leverages the strengths of both Convolutional Neural Networks (CNNs) and attention-based modules. The VAE encoders output the embeddings of Vh and CDR3h of the BCRs in the bottleneck layer, with 15 and 15 neurons, respectively. During training, the VAEs reconstructed the Vh and CDR3h Atchley factor matrices from the bottleneck layers, and compared them against the original Atchley factor matrices, to minimize the loss function.

The antigen embedding part of Cmai is essentially an encapsulation of RoseTTAFold, retaining its original internal structures. In short, antigen sequences are entered into the RoseTTAFold database for multiple sequence alignment (MSA) and template searching. Subsequently, all MSA/templates along with the sequence data are fed into the RoseTTAFold model. The output is derived from the last embedding layer, in the dimensions of (LEN, LEN, 288), where LEN is the length of the antigen protein.

The final binding prediction module of Cmai is built upon the Vh/CDR3h encoders and the antigen encoder. The output embeddings of the Vh and CDR3h encoders are first concatenated, forming a vector of 30 neurons. Then these 30 neurons were inserted at each location of the antigen embedding, forming a tensor of dimension (LEN, LEN, 318). After processing through alternating fully connected layers, average pooling layers and self-attention pooling layers, a logit reflecting the antigen-BCR binding strength is calculated as the final output.

The details of the Cmai model structure have been illustrated in **Fig. S1**.

### Training/testing data creation and model training

To train the VAE embedding model for Vh sequences, we collected a combined dataset of germline Vh sequences from IMGT and Vh sequences observed from real antibodies (“somatic” Vh sequences), with the data described in **Table S1**. To train the VAE model for CDR3h sequences, we collected the CDR3h sequences observed from real antibodies, with the data also described in **Table S1**. The original Vh and CDR3h sequences were fed into the VAE encoders, as well as sequences with random mutations of insertions, deletions, or substitutions, to simulate the somatic hypermutation process. The probability of a sequence undergoing a substitution (with an alternative amino acid randomly picked), deletion, or insertion (with an amino acid randomly picked) at a random locus is equal to 0.25 for each. The probability of staying unchanged is also 0.25.

For training the final binding prediction module of Cmai, we employed a contrastive learning scheme. For each true (positive) BCR-antigen binding pair, we sampled a negative BCR (Vh+CDR3h) with presumably no binding or weaker binding to serve as a contrast. This negative BCR is paired with the same antigen in the positive pair. Then Cmai was trained by inputting both the true positive and the negative BCR-antigen pairs into the model and the loss function requires Cmai to output a smaller logit for the true pair than the negative pair. We collected and created two types of negative BCRs for two phases of our training process: binary training and continuous training. The positive pairs in the binary training data are just formed by known BCR-antigen binding pairs. The negative BCRs are generated by mutating the known binding BCRs randomly three times, via substitutions, deletions and/or insertions (three amino-acid distances in total). We generated a total of 108,888 such (positive BCR, positive antigen)+(randomly mutated negative BCR, positive antigen) pairs. This means that for each positive BCR, there are three corresponding negative BCRs for each paired antigen. In the continuous training data, we focused on a set of our collected pairing datasets that have quantitative measurements of the binding affinity between each positive BCR and antigen pair (such as those from Libra-seq). For each positive BCR-antigen pair, we extracted all BCRs that have no more than 4 amino acid changes from the positive BCR and that have weaker binding affinity compared with the positive BCR, according to this quantitative measurement. Then we formed training data in the format of (positive BCR, positive antigen)+(weaker binding negative BCR, positive antigen). We created a total of 377 such pairs from our data. After the pairing data were created, we split 80% of the data into the training set and 20% of the data into an internal validation set. The training and testing of Cmai on these data were performed for 50 Epochs, where, in each Epoch, Cmai was trained on the binary and continuous training in sequential order.

Cmai was evaluated on the independent test dataset created from pairing data from different and independent studies (described in **Table S1**). For each positive binding pair, we randomly sampled a negative BCR from our library of collected BCR sequences mentioned above, to pair with the same antigen. We sampled a total of 10 negative BCRs, for each positive pair. Then we removed, from these independent test pairs, the BCR-antigen pairs that have appeared before in the training dataset.

More details regarding the training data collection, curation and creation can be found in **File S1** and **Table S1**.

### Cmai antigen binding score calculation

We devised a metric named BCR binding score to represent the overall binding activity of a BCR repertoire towards a specific antigen, based on Cmai predictions. We achieved this by integrating Cmai predictions with the BCR clonal expansion information. Technically, given a sequence of BCR clonotypes in decreasing order of clonal sizes: BCR_1_,BCR_2_,BCR_3_,…,BCR_n_, we first compute, via Cmai, their predicted binding percentile rank%s, r_1_,2_2_,…, r_n_, for a given antigen. Then, based on a series of pre-selected cutoffs Cs (C_1_,C_2_,C_3_,…,C_t_), the percentage of binding BCRs is calculated among the top C_t_ most expanded BCRs. The final binding score is then computed as the mean of all these percentages over the threshold:

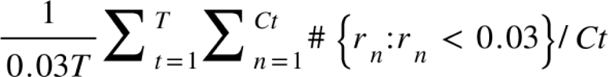

where 0.03 refers to our default rank% threshold to call binding vs. non-binding BCRs. It can be changed to other cutoffs as needed. For bulk RNA-sequencing data-based analyses and most analyses in general, the C cutoffs of 10, 15, 20 were recommended and selected. The C cutoffs for the spatial transcriptomics/BCR data from Engblom *et al*^35^ are selected at 3, 6, 9,12 and the rank% threshold to call binding BCRs was also adjusted to 0.05 instead of 0.03, due to concerns related to the sparsity of the spatial sequencing data.

### Classification of human protein localization

The human_compartment_integrated_full.tsv file is downloaded from: compartments.jensenlab.org/Downloads ^52^. A protein can be located at multiple locations (multiple annotation entries for one protein). We only kept localization annotations with a confidence score>4. Within the remaining annotations, if a protein is documented as an extracellular protein for any entry, this protein would be classified as extracellular. For the remaining proteins, if a protein’s entries are all intracellular, this protein would be classified as intracellular. For proteins that are not classified as either extracellular or intracellular in the above two steps, their localization is recorded as uncertain.

### ICI irAE patient cohort and generation of data

The study was performed following protocols approved by the Institutional Review Board of the University of Texas Southwestern Medical Center (“Biospecimen procurement for immunologic correlates in cancer”, IRB number is 082015-053). Patients provided written, informed consent for additional data collection, presentation, and publication. Clinical, radiographic, and laboratory data were collected from electronic health records (Epic Systems, Verona, Wisconsin). Due to the complexity and difficulty of diagnosing and characterizing irAE, each case was reviewed independently by two clinicians experienced in ICI administration and monitoring, with any differences resolved through third-party adjudication. Patient samples were centrifuged at 3000 rpm at 4°C for 15 min to obtain plasma. Peripheral blood mononuclear cells (PBMCs) were isolated from samples using density gradient centrifugation in Ficoll-Paque Plus Media (Fisher Scientific, Waltham, Massachusetts).

For generation of RNA-seq data, the Agilent 2100 Bioanalyzer system and the RNA Nano and Pico chip kits (catalog # 5067-1511 and 5067-1513) were used for RNA quality measurement. The Illumina TruSeq® Stranded mRNA Library preparation kit (catalog number 20020594) was used for RNA-seq library construction. The library preparation workflow involved purifying the poly-A containing mRNA molecules using oligo-dT attached magnetic beads, mRNA fragmentation, first and second cDNA synthesis, end repair and “A” tailing, ligation of the UMI adapters (manufactured by Integrated DNA Technology), followed by two rounds of AMPure XP beads purification, and then PCR amplification to generate the final cDNA libraries. All libraries must meet the QC requirements before we can proceed with sequencing. All samples were sequenced on the Illumina NovaSeq 6000 sequencer platform with PE-150 sequencing protocol.

For auto-antibody profiling, serum IgG reactivity was tested with a microfluidic antigen array comprising 63 common autoantigens along with two controls (HEL, BSA). These antigens represent common autoantigens implicated in various immune, rheumatic and autoimmune conditions such as SLE50, rheumatoid arthritis, myositis, *etc*. Serially diluted anti-human IgG was added in 4 additional wells to calculate the relative serum IgG levels. The autoantigens were printed in duplicate on nitrocellulose membrane-coated slides. Each slide assayed 15 serum samples plus one PBS control. In brief, 2 ml of serum was diluted 1:100 in PBS with 0.01% Tween 20 and applied to the slide array using microfluidic applications. After washing, IgG isotype antibodies binding to the diverse antigens were detected with Cy-3 conjugated anti-human IgG (1:1000) from Jackson ImmunoResearch Laboratory. Array slides were scanned with GenePix 4400A scanner (Molecular Devices) to generate images for each array. This was converted to Genepix Report file (GPR) using Genepix Pro7.0 software (Molecular Devices).

The averaged fluorescent signal intensity of each antigen was subtracted by local background and the PBS control signal. The net fluorescent intensity of each antigen was normalized by a robust linear model using internal positive controls with different dilutions to generate the normalized fluorescence intensity (NFI) values.

### TCGA data pre-processing

The pre-processed TCGA gene expression data were downloaded from the TCGA Firehose: gdac.broadinstitute.org. To eliminate duplicates, the GBMLGG, COADREAD, and KIPAN cohorts were excluded. There are count matrices for a total of 36 different tumor cohorts. These count matrices were further normalized using total library size, followed by logarithmic transformation (log(x+1)). Subsequently, the transcript-level gene expression data were aggregated to the gene level, resulting in a total of 20,502 genes. The gene expression data for the tumor samples and the normal samples were separated. Genes with average tumor expression level exceeding seven times the average in the normal samples and greater than 0.1 (after library size normalization and log-transformation) were retained as “putative tumor antigens” for each cancer type. The putative tumor antigen lists for all cancer types were then overlapped. The protein sequences encoded by these genes were then mapped in UniProt, which resulted in a total of 1,247 unique putative tumor antigens for all cancer types combined.

### Spatial BCR-seq data preprocessing

The spatial BCR-seq data were acquired from Engblom *et al*^35^. Out of all spatial BCR-seq slides available in the human breast cancer subset, regions c2 and e2 were selected for our analyses based on quality control using UMAP and spatial plot of sequencing spots based on provided cell types. These two slides were the only ones that exhibited reasonably delineated cell type boundaries between the tumor and stromal regions. To identify putative tumor antigens, we started with the tumor antigens for breast cancer in the section above, and further required the expression of the antigens to be higher in the tumor regions than the stromal regions in the spatial BCR-seq datasets. We further narrowed down to extracellular antigens only, which yielded the three tumor antigens presented in **Fig. 3**.

To combat the sparsity and relatively lower quality of the spatial BCR-seq gene expression data, we expanded all antigens’ expression by calculating the spearman correlations of each antigen with all other genes using cells from the tumor region. The top 10 genes (including the putative antigen itself) with correlation coefficients with the largest absolute values were selected. The gene expression of the putative antigen was replaced by the weighted average of these 10 genes, with the weights equal to the correlation coefficients. The expression of the surrogate “gene signature” was used in all downstream analyses, instead of the original single gene’s expression.

### Statistics & reproducibility

Computations were mainly performed in the R (4.2.3) and Python (3.8) programming languages. The Cmai model was trained and validated using Python 3.8, PyTorch version 2.0.0, and pytorch-cuda 11.7. The latest version v1.1.0 of RoseTTAFold as of this paper’s publication was used. All statistical tests were two-sided unless otherwise described. For all boxplots appearing in this article, box boundaries represent interquartile ranges, whiskers extend to the most extreme data point, which is no more than 1.5 times the interquartile range, and the line in the middle of the box represents the median. Enriched pathways under the P-value threshold of 10E-3 and the FDR threshold of 10E-3 were kept. Protein structures were visualized by PyMol 2.5.7 in macOS Sonoma. The circos plot was visualized by ggplot2 (v3.4.4). Kaplan-Meier analyses were performed by the R survival (v3.5.7) package. The R Seurat (v4.4) package was used for processing the spatial BCR-seq data.

## Author contributions

Conceptualization: J.H., T.W.

Data Curation: F.J.F., M.S.V., D.M.Y., J.L., Y.X., C.L., I.R., C.Z., J.E.D., J.H., S.R., S.Y., M.E.G., D.H., Y.G., Y.X., D.E.G. (provided the UTSW irAE cohort), B.S., K.W., T.W.

Formal Analysis: B.S., K.W., S.N., J.Y., Y.G.

Funding Acquisition: T.W., D.E.G., J.H.

Investigation: B.S., K.W., S.N., J.Y., Y.G.

Methodology: B.S., S.N., J.H., T.W.

Visualization: B.S., K.W., S.N.,T.W.

Project administration: T.W. Software: B.S., K.W.,S.N., J.Y.

Supervision: T.W.

Writing – original draft: All authors Writing – review & editing: All authors

## Data and materials availability

The UTSW irAE patient cohort RNA-seq data can be accessed on the Database for Actionable Immunology (DBAI) website (dbai.biohpc.swmed.edu)^4,49,50^ and their clinical characteristics are available at **Table S3**.

The source codes for the Cmai tool are available at github.com/ice4prince/Cmai

## FIGURE AND TABLE LEGENDS

**Table S1** Training and validation datasets.

**Table S2** Classified extracellular *vs.* intracellular proteins.

**Table S3** irAE cohort patient clinical characteristics.

**Table S4** The full list of all auto-antigens used for the irAE analysis.

---

**File S1** BCR/antibody-antigen pairing data curation details.

**Fig. S1.**
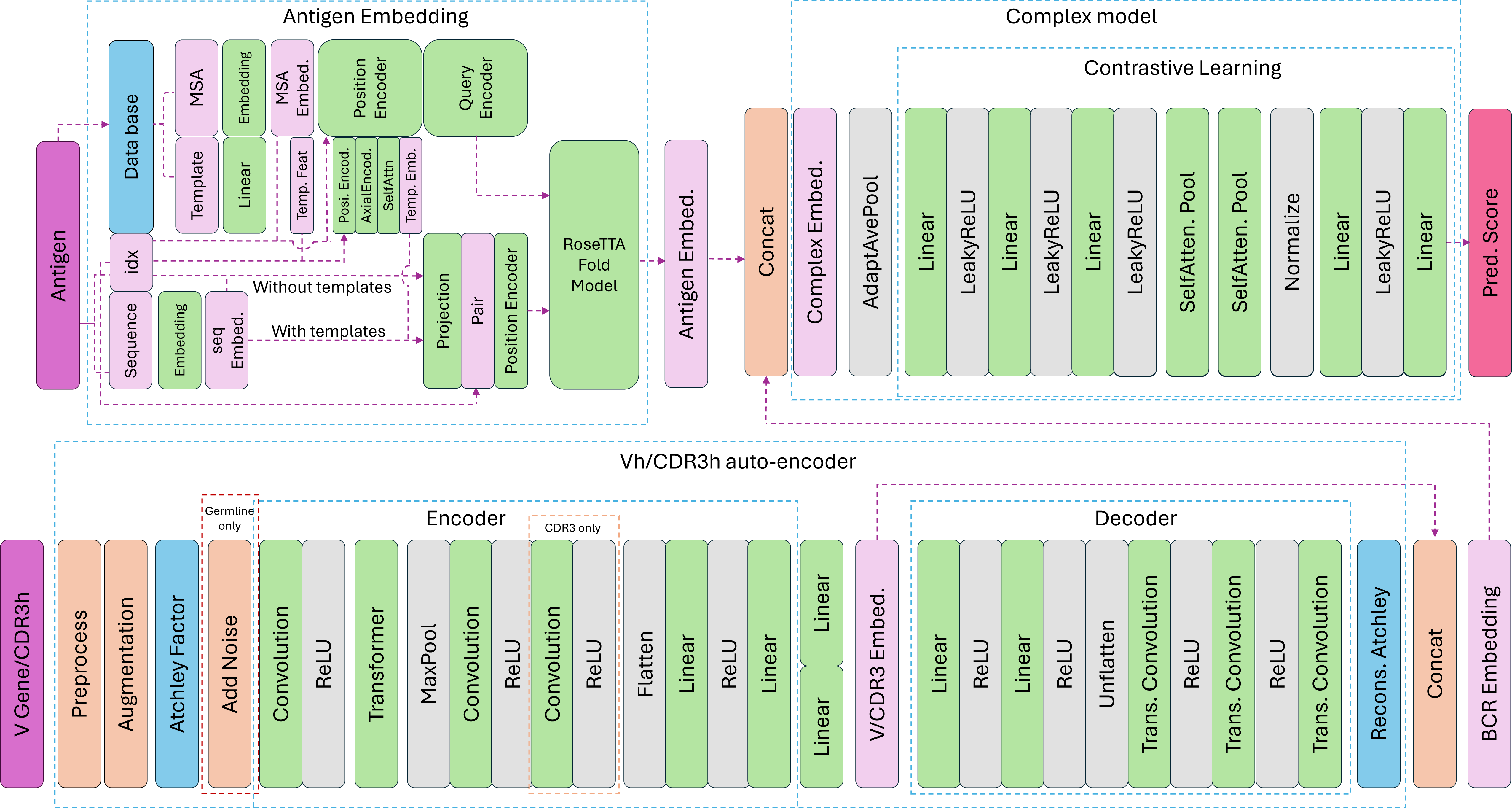
Detailed model structure of Cmai.

**Fig. S2.**
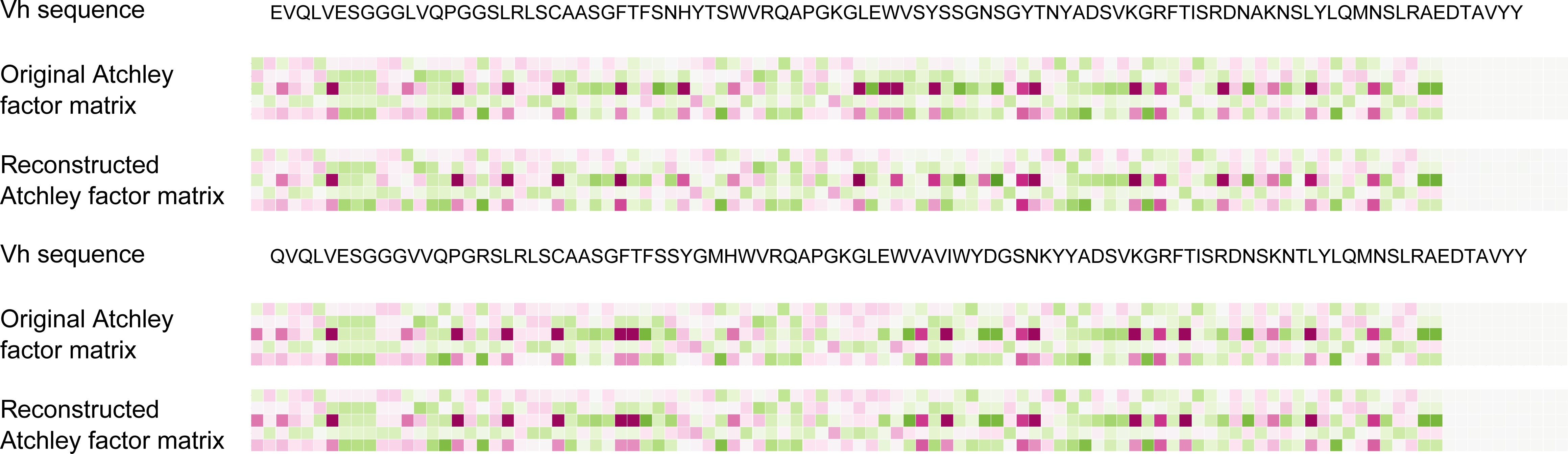
The input and reconstructed Atchley factor matrices of one example BCR Vh sequences.

**Fig. S3.**
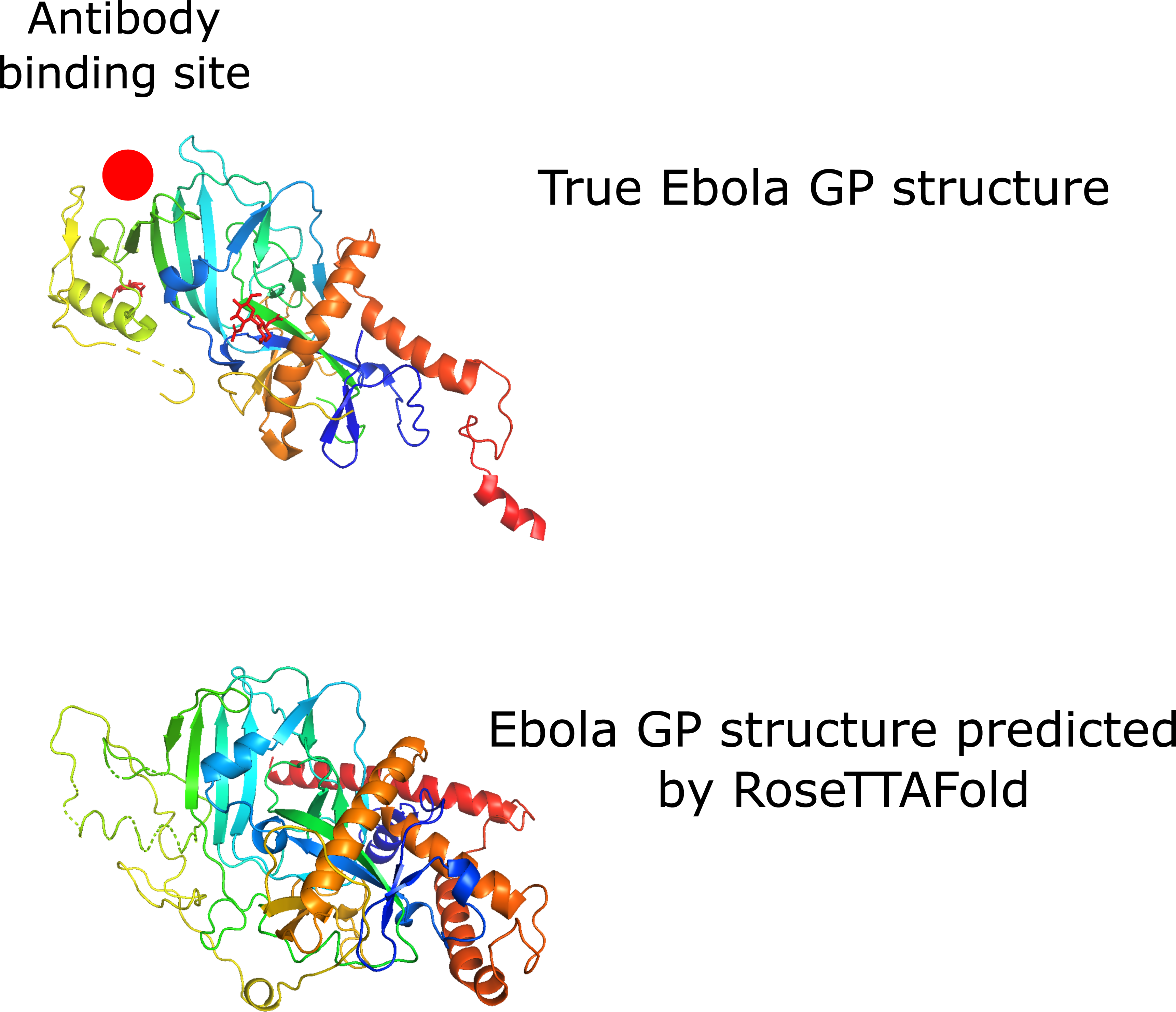
True and predicted Ebola glycoprotein structures. The known antibody binding site is highlighted with a red circle.

**Fig. S4.**
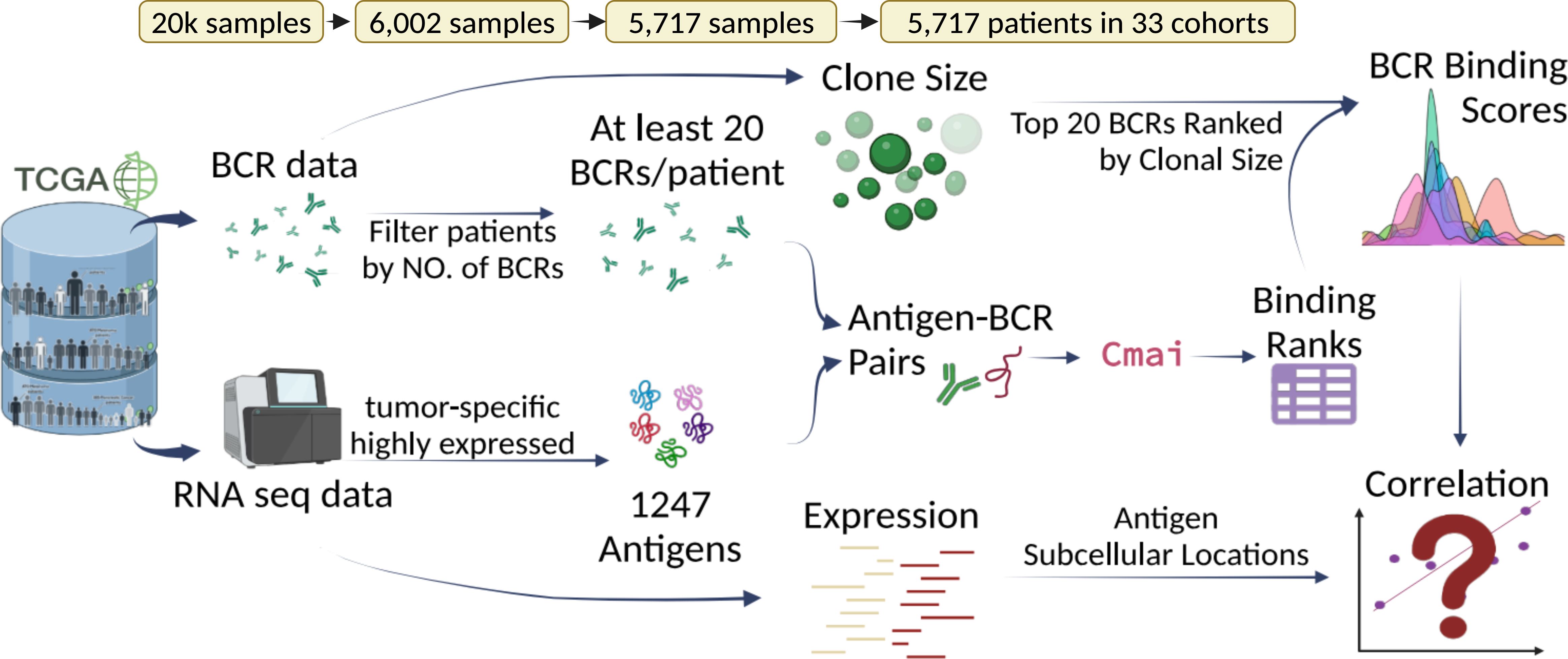
Schematic diagram showcasing how the data from the TCGA were processed and Cmai predictions were made between BCRs and tumor antigens.

**Fig. S5.**
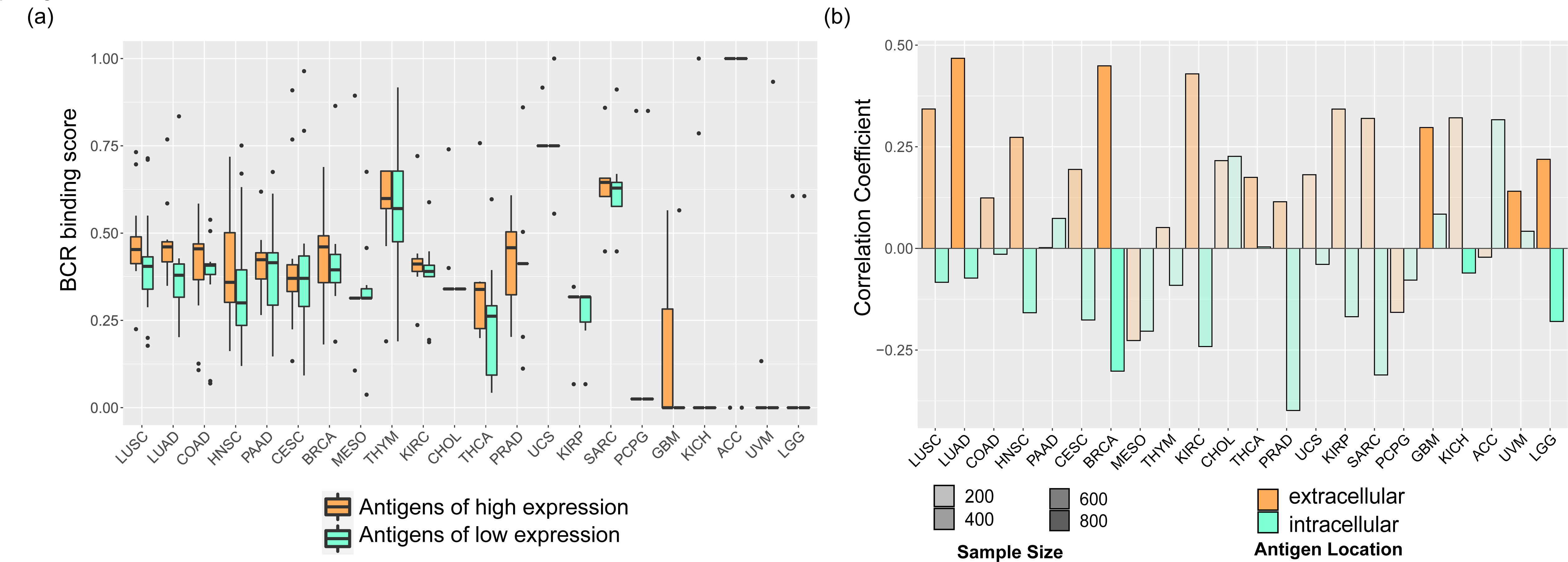
Correlation between tumor antigen expression (average expression > 0.1) and Cmai BCR binding scores. (a) The average BCR binding scores are higher for tumor antigens that are more highly expressed, in each TCGA cohort. (b) The correlation between BCR binding scores and tumor antigen expression is higher for extracellular antigens than for intracellular antigens.

**Fig. S6.**
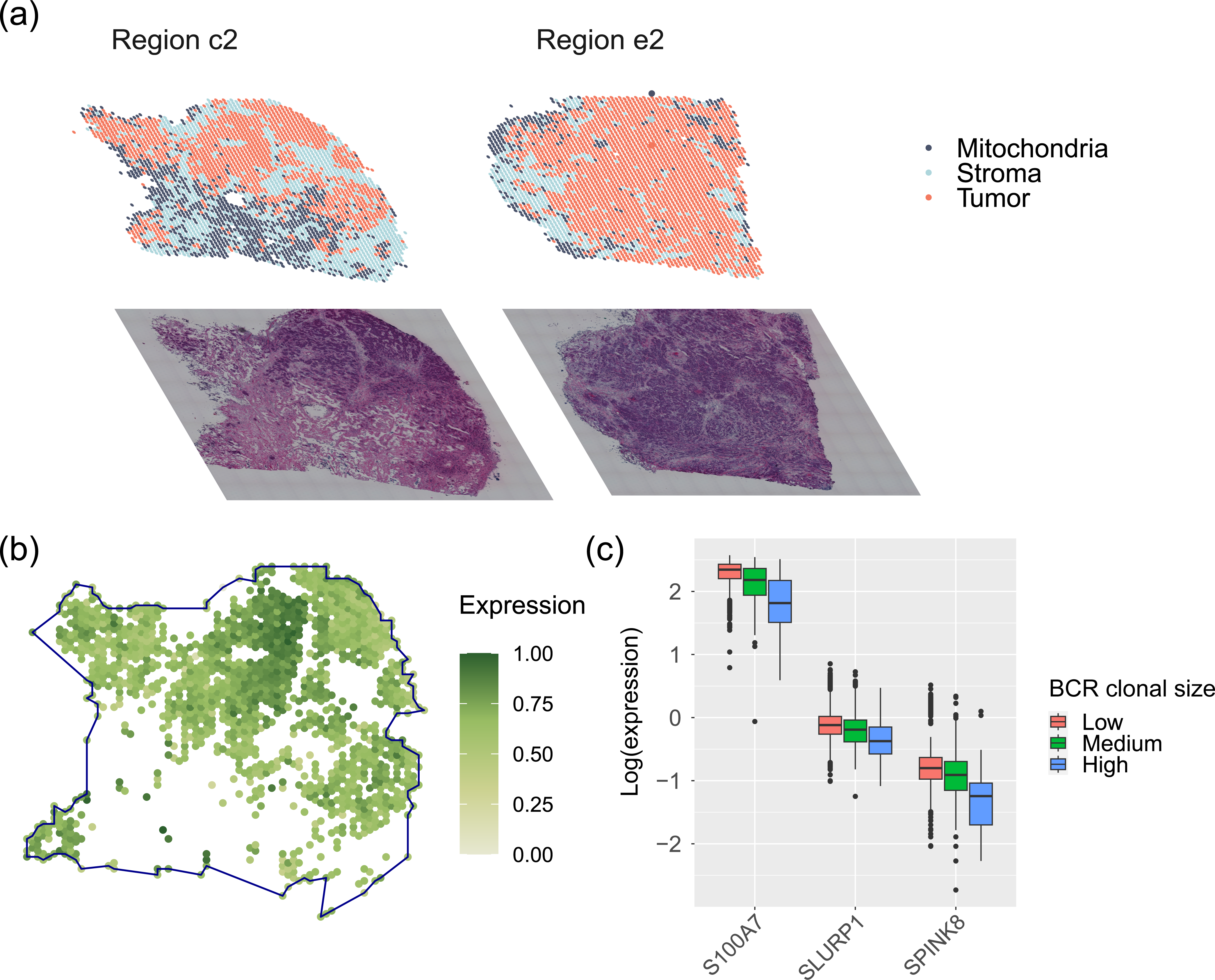
Cmai reveals BCR-dependent co-localization of B cells and tumor cells. **(a)** The spatial distribution of the spatial sequencing spots, overlaid with the corresponding H&E staining images. (b) The spatial expression pattern of *S100A7* in the tumor sequencing spots. (c) Tumor antigen expression in the tumor sequencing spots of the c2 region with high, medium and low BCR clonal expansion sizes.

**Fig. S7.**
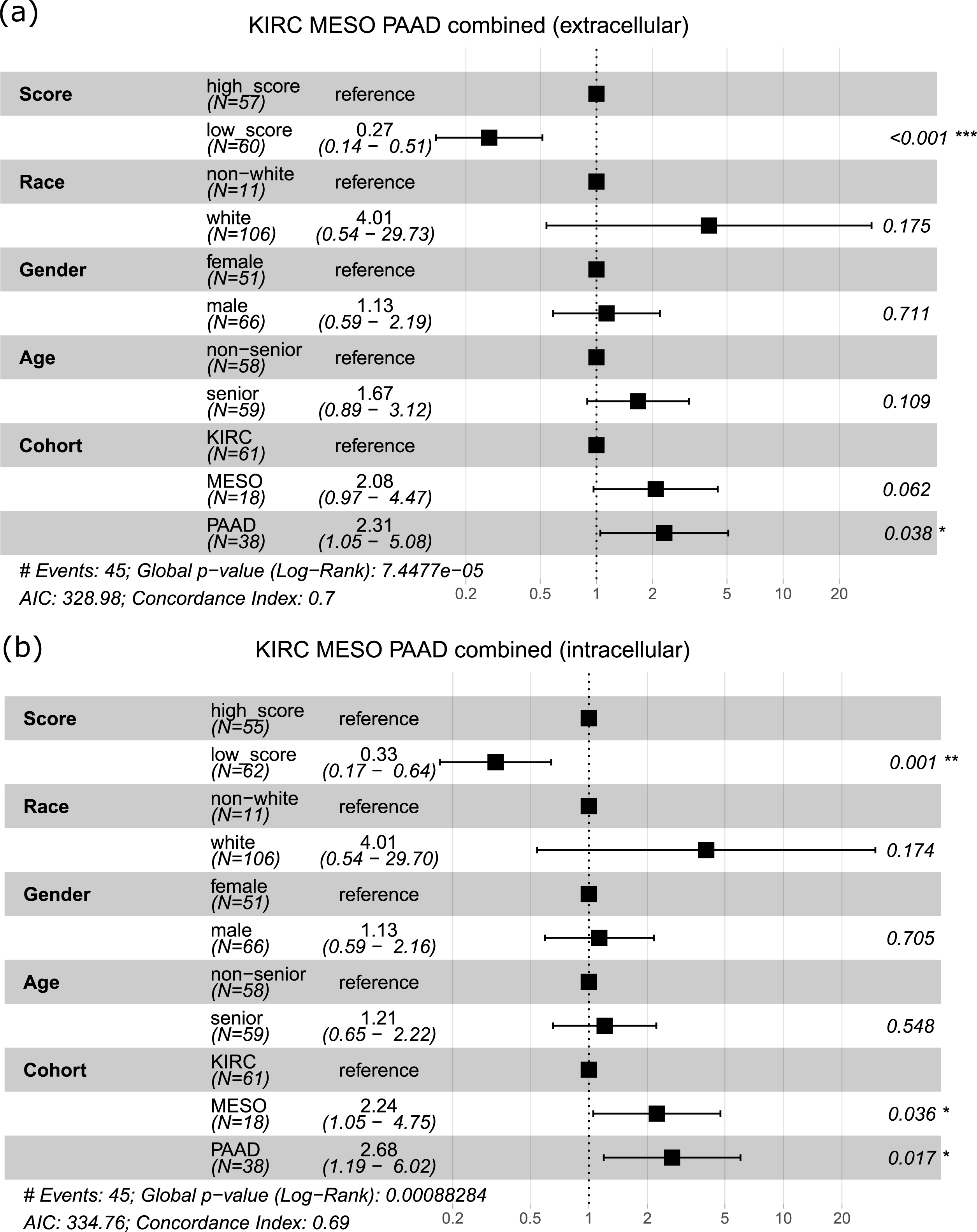
Forest plot showing the results of the multivariate correction for the prognostic effects of the Cmai BCR binding scores, in the KIRC, MESO and PAAD patients combined. (a) Binding scores calculated for extracellular antigens only; (b) binding scores calculated for intracellular antigens only.

**Fig. S8.**
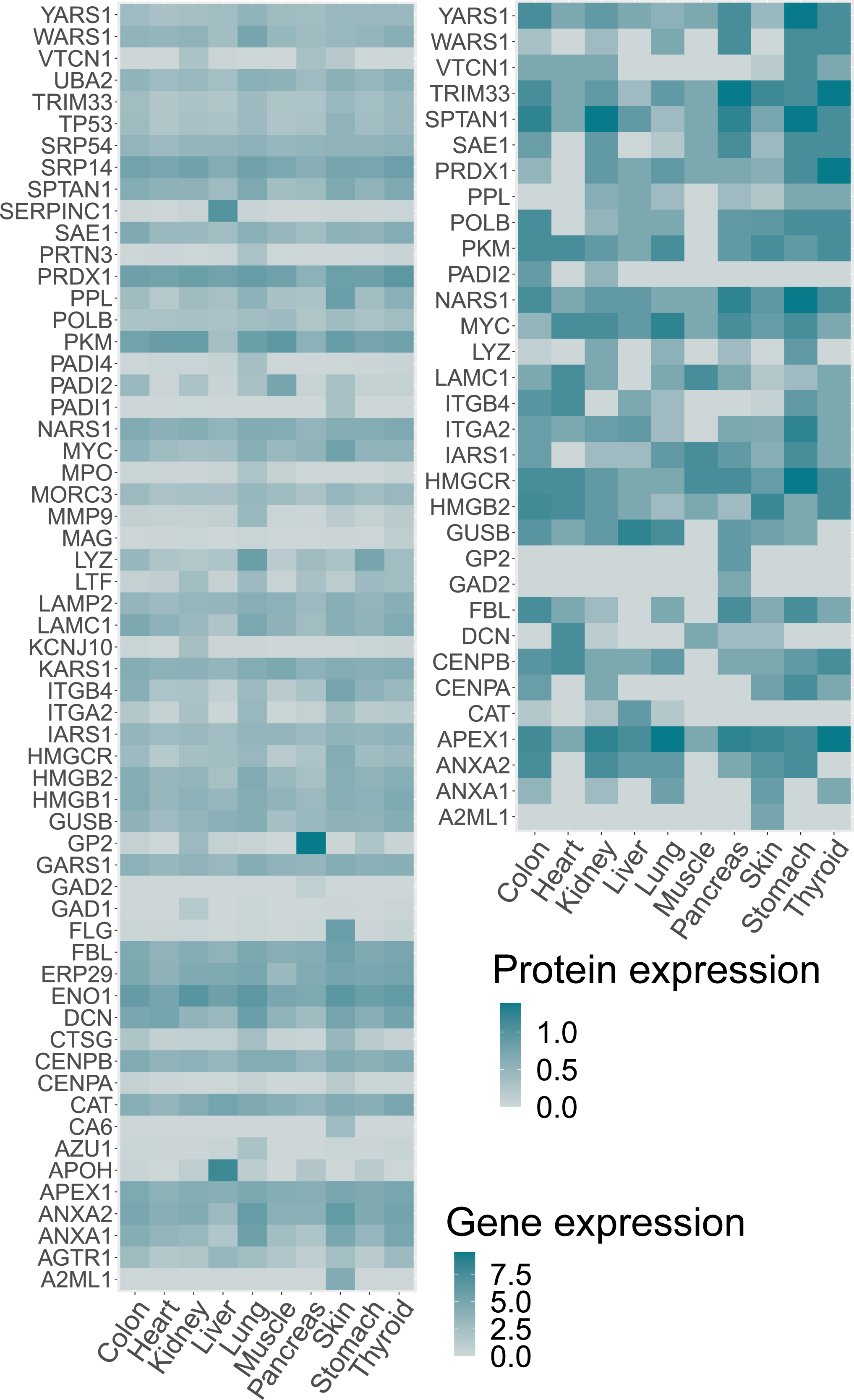
The RNA and protein expression of tumor antigens in the organs affected by the irAEs captured in our patient cohort.

