## Supplementary material for "Cmai: Predicting Antigen-Antibody Interactions from Massive Sequencing Data": Sup. File 1

### Sup. File 1: BCR/antibody-antigen pairing data curation details

#### Dataset selection

We searched for published literature (including databases such as IEDB) that reported antibody/BCR-antigen pairing data. These datasets include pairing data from a variety of sources, such as neutralization assays or high-throughput sequencing methods such as Libra-seq. We focused on the heavy chains in this study and only kept records for which the heavy chain sequences were available. We split our datasets into training and validation based on consideration of sample sizes and sufficient variety of antigens in the both cohorts. In the context of this particular study, BCRs and antibodies are two exchangeable concepts, and thus we do not distinguish between them. The full list of the selected datasets were provided in **Table S1**.

For all BCRs provided in each cohort, IgBLAST [1] was used to parse the protein or nucleotide sequences. We used the efficient PyIR[2] implementation for all nucleotide sequences, whereas the IgBLAST web tool was used for protein sequences. We split the BCR heavy chain protein sequences into two parts as two different inputs of our machine learning model. One part ("CDR3h") contains the CDR3H sequence, starting from C and ending with W, and the other part ("Vh") contains the segment of the somatically mutated BCR that is upstream of CDR3Hs, excluding the C. On the other hand, we curated antigen protein sequences for all our BCR-antigen pairing records as another input of our model. We filtered out pairing records with antigen, Vh or CDR3h entries containing amino acids marked as "X" or "*". Less than 0.03% were removed at this step.

#### Binary and continuous training data

The Cmai model is trained based on the idea of contrastive learning, where we always compare between a positive antigen-binding BCR with better binding affinity and a negative BCR with worse or no binding affinity against the same antigen. The data we collected in the above step mostly contain positive binding pairs. According to the nature of the collected data, we created two types of negative binding pairs, for the contrastive learning schema.

For the majority of the data, we know a BCR is binding to an antigen, which is binary and no further quantitative measurements of the binding affinities are provided. In such cases, we created randomly mutated BCRs based on the positive binding BCRs to generate negative BCRs that are very likely to have weakened binding and that still have similar protein sequences to the positive binding BCRs. This approach is also in the spirit of diffusion models where noises are added to the positive training cases and the model is asked to discern the noises. On the other hand, for some of the training data that we collected, there is a quantitative measure (such as Libra-seq score) for quantitatively assessing the binding affinities of the BCRs against an antigen. In such cases, we derived matched records of two BCRs with similar protein sequences and different binding affinity against a certain antigen, according to the quantitative score. Details of the data generation processes are described below.

Generation of the binary training data: The *IEDB*, *CoV-AbDab*, *Lee*, *Guthmiller*, *Chen*, *Sutton*, *Kubota*, *Xiao*, and *Peissert* cohorts were used to generate binary training datasets. The details and citations for each cohort are included in **Table S1**. For each antigen-antibody pair, binding Vh and CDR3h sequences of the binding BCRs were previously identified using the preprocessing steps mentioned above. These binding BCRs are the positive "binding" BCRs in the contrastive learning scenario. Then we employed random mutations to simulate the negative BCRs for each positive BCR-antigen pairing record, with the simulation procedure designed in respect of our known knowledge of the pattern of BCR somatic hypermutation.

For each negative BCR, we created a total of 3 mutations, which likely has partially or fully abolished the binding capability of the original positive BCR towards the target antigen. To account for the differences between framework regions (FWR) and CDR regions, we set the probability of mutations occurring in each region to the following: $FR1 \approx 0.13$, $CDR1 \approx 0.28$, $FR2 \approx 0.13$,$CDR2 \approx 0.21$, $FR3 \approx 0.10$, and $CDR3 \approx 0.15$ according to rough estimates of the mutation frequency data presented in previous reports [3,4]. Given its relatively short length, we further ensured that CDR3 has at least one mutation. We sampled the locations of mutations using indices of existing amino acids. After 3 locations were selected, we sampled mutation types. The possible types of mutations consist of substitutions, deletions and insertions. We set the probability of occurrence of each of these types of mutations to 0.9, 0.05, and 0.05, according to rough estimates from previous reports [5,6]. For substitutions, the existing amino acid was replaced by a random sampling of the remaining 19 amino acid choices; for deletions, the selected amino acid was simply deleted; for insertion, we randomly sampled from the total of 20 amino acid choices. This negative BCR sampling was repeated 3 times for each binding BCR, to create three mutations.

Generation of the continuous training data: The *Libra-seq*, *Shiakolas*, *Dugan*, *54042-4*, and *46472* cohorts were sources for the continuous training data. In each cohort, a binding affinity score (*e.g.* Libra-seq score, UMI count, ic50 value, *etc.*) for each antigen-antibody pair was available. We identified a cutoff value for each cohort (usually according to suggestions of the original publication) to define binding and non-binding BCRs. Then we generated all possible pairs of BCRs with <4 edit distances (smaller than 4 amino acid differences in Vh and CDR3h combined) for all cohorts. For each BCR pair, the two BCRs additionally need to satisfy (1) the better BCR is a binding BCR as defined above, and (2) the worse BCR has a worse binding score than the better one for the same antigen, no matter binding or non-binding.

### Validation data

The *Mason*, *Shan*, *Chappert*, *Makowski*, and *Hie* cohorts were curated for the validation of Cmai. For these cohorts, we matched positive binding BCRs with randomly sampled BCRs from our background BCR pool as the negative cases.
